# Distinct cortex-wide dynamics underlie evidence accumulation, short-term memory and choice

**DOI:** 10.1101/2025.07.24.666672

**Authors:** Renan M. Costa, Jiaqi K. Luo, Peter S. Salvino, Lyn A. Ackert-Smith, Shawna M. Ibarra, Lucas Pinto

## Abstract

Most decisions involve multiple cognitive processes. Recent findings suggest that these processes are distributed across the cortex, but how each of them is distinctly implemented remains unknown. Here, we designed a virtual-navigation task for mice that dissociates the accumulation and short-term memory of sensory evidence, and choice. Combining decoding models with cortex-wide widefield Ca^2+^ imaging, we observed distributed and dynamic encoding of the task’s decision variables both within and across areas. Further, the time course of the decision variable as it was accrued and then held in memory was well predicted by an area’s spontaneous timescales measured before learning, suggesting that these computations co-opt intrinsic circuit properties. Finally, we show that the dynamics underlying different cognitive processes evolve in orthogonal subspaces, and that this geometry breaks down during erroneous choices. Thus, we reconcile previous findings by showing that cortex-wide dynamics supporting distinct cognitive processes are disentangled.

## Main

Decision-making tasks engage neural activity across the cerebral cortex of rodents and primates, with activity within each cortical area simultaneously reflecting multiple cognitive, sensory and motor processes^1–21^. Moreover, different areas of the mouse cortex display surprisingly similar coding properties during perceptual decisions^5–7,9,16,21–23^. Taken together, these findings may suggest that large-scale cortical dynamics supporting different cognitive processes are entangled—i.e., each process is similarly distributed across the cortical sheet, with little functional specialization. Alternatively, they could be implemented by distinct (time-varying) activity patterns across partially overlapping large-scale networks, as suggested by neuroimaging studies in humans (e.g., ^11,24–26^).

This issue has remained unresolved in part because most behavioral-task designs are not conducive to adjudicating these competing hypotheses. For example, in many tasks, the process of motor-choice formation correlates, and temporally overlaps with, the gradual accumulation of sensory evidence^27–29^. This means that neural activity cannot be unequivocally attributed to each process. Furthermore, evidence accumulation itself is a working-memory process^30^, as it involves both memory of the evidence tally and updating it with new incoming evidence. However, it is presently unclear if its underlying neural dynamics are shared with other types of working memory, e.g., in which a stable memory needs to be maintained and then manipulated towards a decision. To overcome these limitations, we developed a new behavioral task for mice navigating in virtual reality (VR), which temporally dissociates three core cognitive processes: gradual accumulation and short-term memory of sensory evidence, and choice formation. We combined this task with widefield Ca^2+^ imaging to simultaneously measure excitatory activity across many cortical areas. Using decoding approaches, we show that each of these processes is encoded dynamically but in orthogonal subspaces, defined by single-area or cortex-wide activity. Thus, while cognitive processes are not strictly functionally localized, they still rely on functionally specialized large-scale networks across the mouse cortex.

### Delayed match to evidence: a new mouse VR task to dissociate cognitive computations

We developed a task building on previous delayed match-to-sample designs for mice, in which they are first presented with a sample stimulus and, after a delay, compare it with one of two test options^19,31–34^. Crucially, in our case the sample is noisy, such that the mice need to gradually accumulate evidence to determine its category (**Fig. 1a, Extended Data Fig. 1, Supplementary Tables 1–4**). As mice navigate a T-maze, they first encounter a bilaterally symmetrical checkerboard along maze walls with black and white squares in different proportions^35^ (1 m; only a 20-cm sliver is visible at a time), and must determine if the checkerboard is predominantly black or white. They then enter a delay region with neutral wallpaper (50 or 75 cm), where they must hold the predominant color in short-term memory. As they approach the end of the maze, we reveal two test stimuli, one black and one white, each along a wall and on randomized sides trial to trial (75 cm). The mice are rewarded for turning into the arm on the side with the test stimulus whose color matches the predominant color of the sample checkerboard. Thus, because they cannot make an informed choice before the test stimuli are revealed, the task dissociates evidence accumulation from motor choice. Further, although working memory is required throughout, because the test and sample stimuli are separated by a delay, the task isolates a period when sensory memory is not updated via accrual or used for a decision. To our knowledge, this is the first mouse task that accomplishes all these dissociations (though note that other processes like reward expectation and decision confidence are not isolated).

**Fig. 1.**
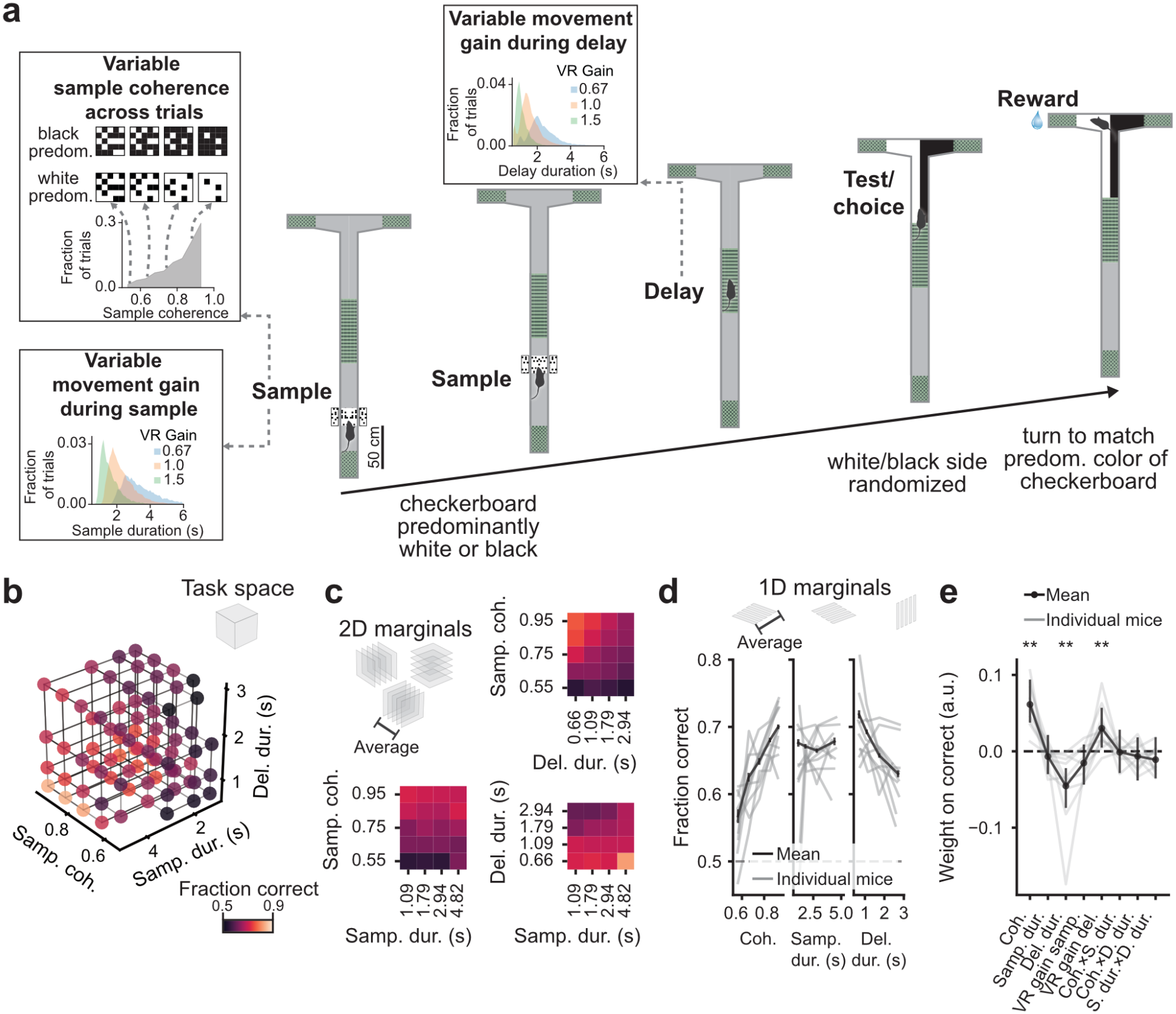
Delayed match to evidence: a new mouse VR task that temporally dissociates cognitive processes. **a**, Schematics showing the progression of an example trial of the task with a predominantly white checkerboard. Insets show the continuous distribution from which checkerboard coherence is drawn, and the partially overlapping distributions for sample and delay duration, resulting from the combination of gain manipulations and running speed distributions across trials. The partial overlap in durations allows us to dissociate the contributions of gain and duration to behavior and neural activity. **b**, Aggregate performance as a function of the combination of three binned task parameters, tiling the 3D task space (*N* = 10 mice, 454 sessions, 49800 trials; averages shown are across all trials). Bins with fewer than 20 trials were removed from the analysis. **c**, Same as panel b, but marginalized along the different 2D slices of the 3D space. **d**, Same as panels b and c, but marginalized along each axis. Gray lines: mice. Black lines: mean of aggregate trial data. Error bars: ± standard error of the mean (s.e.m.) across trials (*n* = 49800). **e**, Coefficients of a logistic regression model fit to predict the probability of correct choice from several task parameters (cross-validated prediction accuracy of 0.60 ± 0.01, mean ± s.e.m.). Gray lines: individual mice (*N* = 10). Black data points and lines: average across mice. Error bars: 95% confidence intervals (CI) bootstrapped per mouse (*n* = 786–7156 trials per mouse), then averaged across mice (*N* = 10). **: *P* < 0.01, Wilcoxon signed-rank test against zero with false discovery rate (FDR) correction.

Another key feature of our new task is that we systematically and independently manipulate several of its parameters (**Fig. 1a** insets). First, we vary perceptual difficulty on each trial by changing the sample checkerboard’s coherence, defined as the relative proportion of squares in the predominant color. Moreover, we vary the duration of evidence accumulation and short-term memory by changing the gain between the mice’s physical running on the ball and their progression through the virtual environment, either speeding it up or slowing it down. Thus, we generate a continuous three-dimensional (3D) space of cognitive processes. Crucially, this allowed us to characterize how cortex-wide dynamics change along each cognitive axis, and the relationship between these axes.

We found that behavioral performance varied systematically within the 3D task space, reaching up to 81% correct with easier trial combinations (**Fig. 1b, c, Extended Data Fig. 2a, b**, *N* = 10 mice, 454 sessions, 49800 trials), with little side, color, or trial-history bias (**Extended Data Fig. 2c–e**). Overall performance across all difficulties was 67%, compatible with working-memory tasks previously developed for mice (e.g., ^19,28^). On average, performance increased with increasing sample coherences, decreased with longer memory-delay durations, and changed little with sample duration (**Fig. 1d**). To jointly quantify how much each task parameter affected behavioral performance, as well as the impact of their interactions, we used logistic regression to predict the mice’s choices on each trial (**Fig. 1e**). Sample coherence and delay duration had significant (*P* < 0.01, Wilcoxon signed-rank test) and similar-magnitude impact on behavioral performance, albeit in opposite directions. The effect of delay duration on performance was not a confound of slower running speeds correlating with task disengagement, since speed negatively predicted correct performance (**Extended Data Fig. 2f**). On the other hand, on aggregate, evidence-accumulation duration did not predict higher performance. This is likely because even the shortest duration in our case is longer than the typical integration time window over which evidence-accumulation performance improves (∼0.5 s)^27,29^. Indeed, model-free analyses showed that only the hardest coherence level benefitted from more prolonged accumulation (**Extended Data Fig. 2a**). Interestingly, task parameters combined linearly to affect performance (*P* > 0.05 for all interaction terms). Finally, we asked how much the mice weighted evidence from different parts of the checkerboard by predicting their choices from the effective coherence at different maze positions using logistic regression. Compatible with a previous pulse-based evidence accumulation task^28^, we observed a variety of individual evidence-weighting profiles. On average, however, the animals took into account evidence from throughout the checkerboard (**Extended Data Fig. 2g**, *P* < 0.01 for all position bins, Wilcoxon signed-rank test).

### All the task’s cognitive processes engage distributed cortical activity

Next, we asked what cortex-wide activity patterns were related to the task’s different cognitive processes. We transgenically expressed the Ca^2+^ indicator GCaMP6s in cortical layer 2/3 excitatory neurons^36^ (**Fig. 2a**) and used a widefield macroscope to image through the intact cleared skull from across the dorsal cortex with mesoscale spatial resolution (∼50 µm/pixel), while mice performed the task (**Fig. 2b, c**, *n* = 5411 trials across 29 sessions from 7 mice). We first averaged activity within different portions of the maze, each corresponding to a different combination of cognitive processes. This revealed partially distinct patterns of multi-area cortical activity for each task epoch (**Fig. 2d**). For instance, while posterior regions such as the visual, posterior parietal and retrosplenial cortices were maximally engaged during evidence accumulation, activity during the memory delay ramped up in frontal regions but decayed in visual ones (**Fig. 2e**).

**Fig 2.**
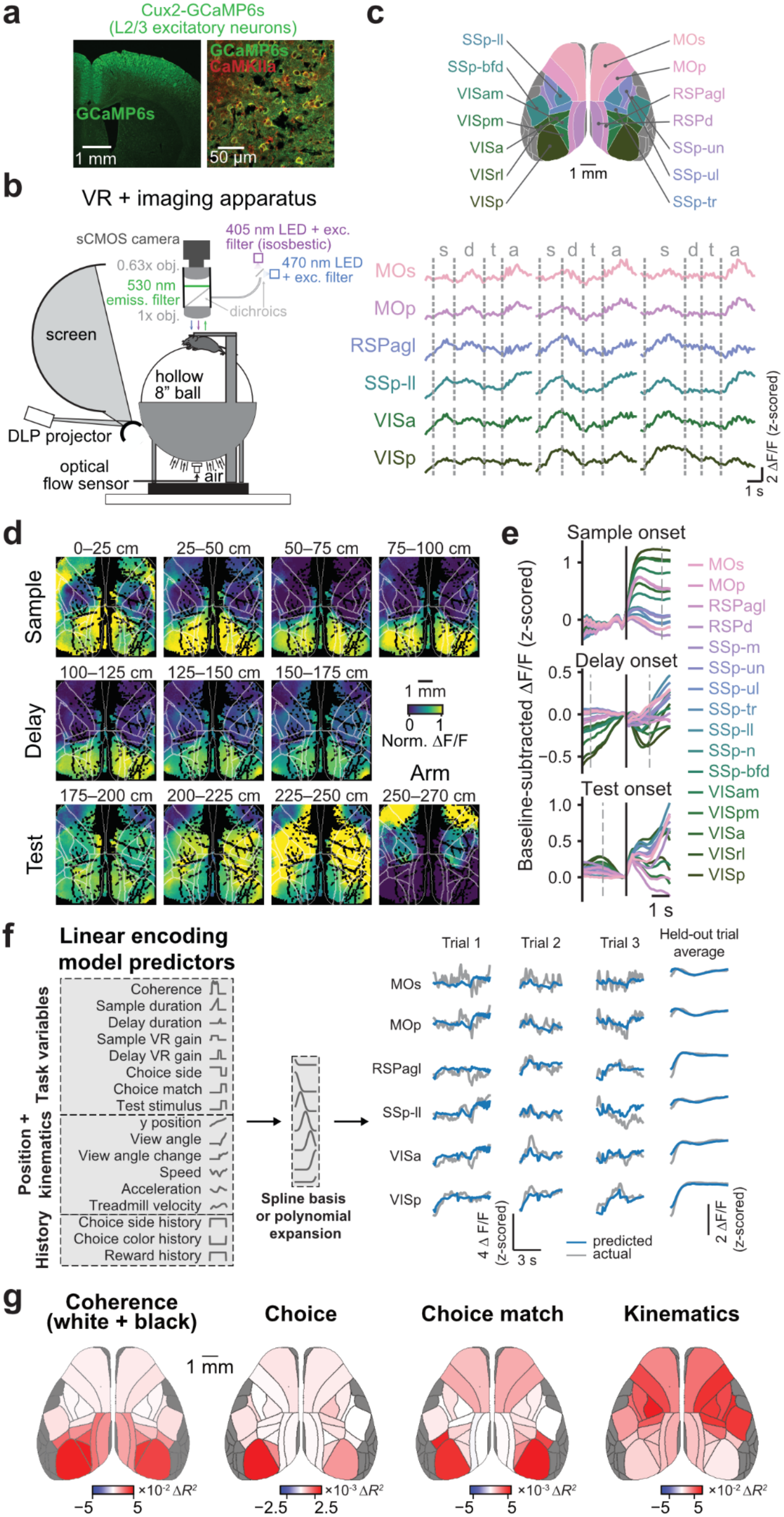
Task variables are encoded by partially overlapping multi-area networks across the dorsal cortex. **a**, Left: confocal image of a coronal brain section, showing GCaMP6s in superficial cortical layers (amplified with GFP immunostaining). Right: immunostaining for GFP (green) and CaMKIIα (red), showing high penetrance and specificity of GCaMP6s expression in layer 2/3 excitatory neurons. **b**, Schematics of mesoscale widefield imaging setup integrated with VR. **c**, Single-trial examples of ΔF/F averaged across pixels within anatomically defined cortical regions (reference map on top). Vertical dashed lines: task epoch onsets; s: sample, d: delay, t: test, a: arm. **d**, Pixel-wise activity across the dorsal cortex for an example behavioral session, averaged for each maze position over correct trials. The activity of each pixel is normalized between its minimum and maximum to emphasize its timing relative to other pixels. White contours outline the cortical regions according to the map in C. **e**, Grand average of ΔF/F for each anatomical region across all mice and sessions (*n* = 3685 correct trials from 29 sessions and 7 mice), aligned in time by the onset of each task epoch (vertical black line). Dashed vertical gray lines indicate median onset time of other epochs. Traces are subtracted by activity during a 0.2-s baseline preceding epoch onset. Error estimates are omitted for legibility. **f**, Schematics of a mixed-effects, L2-regularized linear encoding model of neural activity and example actual and predicted activity traces for individual and averaged left-out trials. **g**, Cortical-area maps for the change in variance explained when models are refitted without sets of predictors, for four example sets. Each model was fitted to data from all sessions (*n* = 4856 trials from 29 sessions and 7 mice), and cross-validated *R*^*2*^ was averaged across five folds. See **Extended Data Fig. 3b** for all predictor sets and **Supplementary Table 5** for statistics.

Next, taking advantage of their partial-to-full behavioral dissociation, we quantified the relative contributions of cognitive, sensory, and motor processes (and their duration) to the activity of each cortical area as a whole, using a linear encoding model with multiple behavioral variables as predictors (**Fig. 2f**). The model predicted held-out data with high accuracy (**Fig. 2f, Extended Data Fig. 3a**). We then dropped different sets of model predictors and refitted reduced models, quantifying their contributions to cortical activity by measuring the change in how much variance the reduced models explained relative to the full one. Most sets of predictors contributed significantly to the encoding model of each area, although to different extents (**Fig. 2g, Extended Data Fig. 3b, Supplementary Table 5**). For instance, while the removal of stimulus- and choice-related predictors affected model accuracy more in posterior than frontal regions, the opposite was true for running-kinematic variables. Memory-delay duration, on the other hand, more evenly explained neural activity across areas. Note that the strong contribution of choice information to posterior cortical activity is compatible with the existence of choice signals in single visual and posterior parietal neurons and their populations in rodents and primates (e.g., ^21,37,38^). It could additionally be related to the fact that our area-level models average over finer-spatial-scale information in frontal areas, as our decoding analyses below suggest. Altogether, our results so far show that dynamics in all of the mouse dorsal cortex reflect evidence accumulation, short-term memory and choice, as well as non-cognitive variables, although the areas differ quantitatively in terms of how much variance each process accounts for. This is largely compatible with previous reports^6,7,9,16,21^.

### Cognitive processes are implemented in dynamic activity subspaces

We next asked whether the distributed cognitive processes we observed unfolded over static or dynamic neural-activity subspaces. We first used principal components analysis (PCA) to decompose pixel-wise activity (a ∼12,000-dimensional space) and visualize how it evolved during each portion of the maze along the first 3 principal components (PCs). Interestingly, these activity trajectories changed directions when task epochs changed, e.g. from evidence accumulation to short-term memory, compared to shallower-curvature trajectories within each epoch (**Fig. 3a**). To quantify this, we computed the angle between trajectory segments within the space spanned by the first 20 PCs (which captured 71.2 ± 2.4 % of the variance, **Extended Data Fig. 4a**). Indeed, we found that the angles between segments within a task epoch were significantly smaller than across epochs (**Fig. 3b**, *P* < 0.001, one-way repeated-measures ANOVA). This was not trivially related to changes in the visual scene and running patterns associated with different maze epochs, since regressing out luminance and kinematic-related changes in neural activity yielded similar results (**Extended Data Fig. 4b–h**).

**Fig 3.**
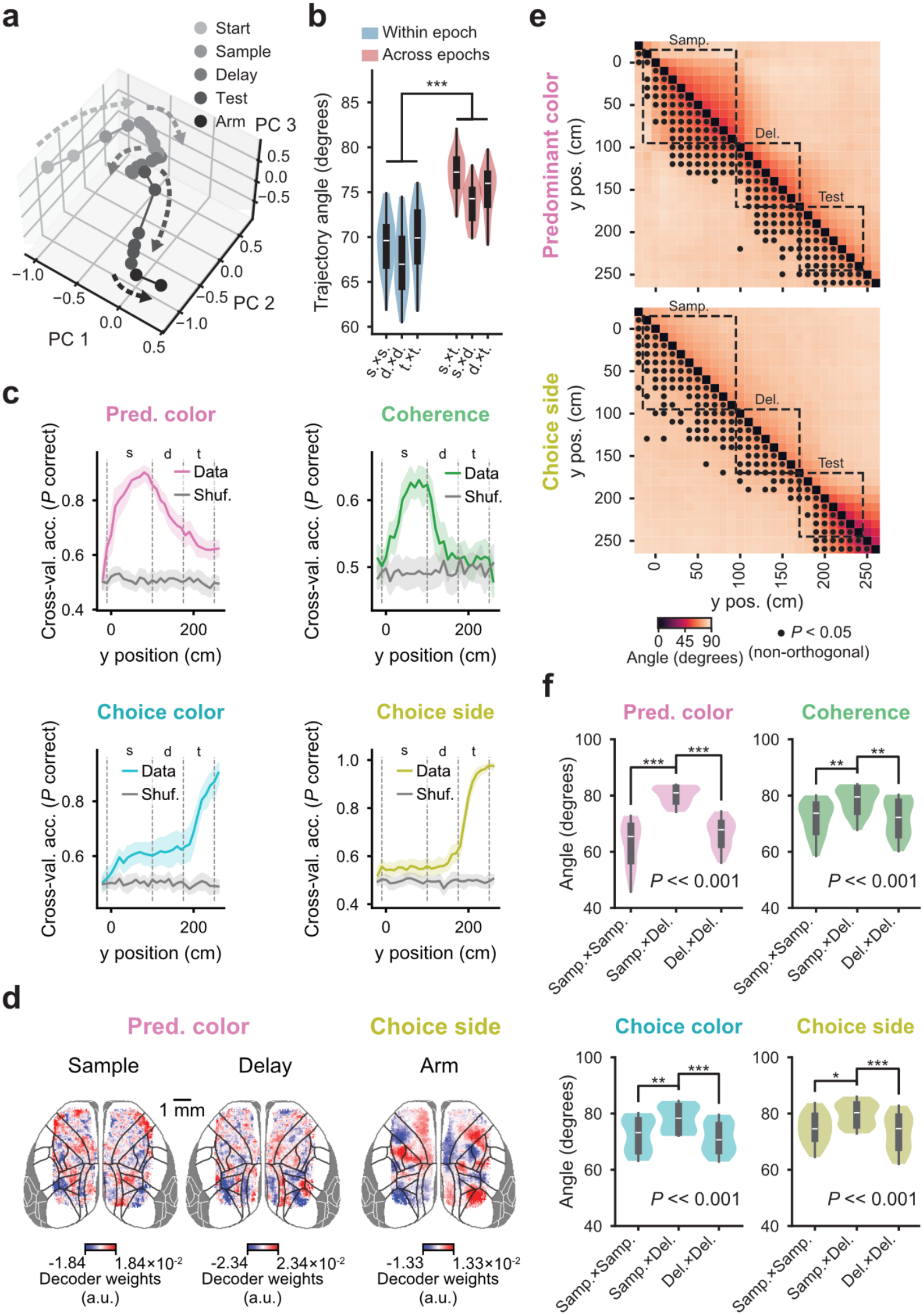
Cognitive processes are implemented in dynamic coding subspaces. **a**, Single-session example of a trial-averaged activity trajectory projected onto the first 3 PCs, computed on the full pixel space. Activity is binned by y position in the maze, with different shades of gray indicating different maze regions. **b**, Distribution of angles between activity-trajectory segments within (blue) and across (red) epochs, projected onto the top 20 PCs. ***: *P* < 0.001 for all individual comparisons between within- and across-epoch angles, Tukey’s post-hoc after one-way repeated-measures ANOVA (*n* = 20 sessions, 5 mice). s: sample, d: delay, t: test. **c**, Accuracy of binary decoders of task variables from the activity of pixels from across the dorsal cortex, assessed on held-out trials. Different decoders were trained for each session and y position (*n* = 20 sessions, 5 mice). 67–315 trials per session were split into training sets (*n* = 53–252 trials per set) and test sets (*n* = 14–63 trials per set). Error shades: 95% CI across sessions (*n* = 20). s: sample, d: delay, t: test. **d**, Distributions of decoder weights across the dorsal cortex, for example task variables and epochs, averaged across 20 sessions. **e**, Angles between the decoding axes for the same task variable, for pairs of maze positions. Dashed boxes: task epochs. Circles: significantly non-orthogonal angles (*P* < 0.05, shuffle test, see Methods for details). Angles and *P*-values are averages across 20 sessions. **f**, Distributions of decoding angles within and across task epochs, for different behavioral variables (*n* = 20 sessions). Printed *P*-value is for a one-way repeated-measures ANOVA with factor ‘epoch pairs’. *: *P* < 0.05, **: *P* < 0.01, ***: *P* < 0.001, Tukey’s post-hoc test. Note that, for all variables, angles are significantly more orthogonal across than within epochs.

Note, however, that the highest-variance activity modes in the data that are found by PCA are not necessarily aligned with the computations in the task (e.g., ^31,39^). Thus, we next sought to determine how large-scale cortical coding of the task’s specific cognitive processes evolved over time, focusing on the main decision variables. We used support-vector machine (SVM) decoders of binarized task variables—’predominant color’ of the sample (i.e., sample category), ‘sample coherence’, ‘choice color’, and ‘choice side’—fitted separately for different maze positions. Specifically, we used the per-session activity of simultaneously imaged pixels from across the dorsal cortex to classify these behavioral variables, learning a weight for each pixel (spatially binned to ∼200 µm/pixel to decrease the number of model parameters and avoid overfitting). These cortex-wide decoders predicted held-out trials with high accuracy, peaking during the appropriate task epoch for each variable (**Fig. 3c**). For instance, accuracy for the ‘sample coherence’ and ‘predominant color’ decoders was maximal during presentation of the checkerboard, although it was smaller in magnitude for coherence than category. Interestingly, unlike coherence, category decodability persisted into the test region of the maze, when the sample category is compared to the test stimulus. On the other hand, decodability was not trivially related to the maze region, as most task variables could be decoded during the memory delay (**Fig. 3c, Extended Data Fig. 5a**). This suggests that information about different cognitive processes co-exists at the level of cortex-wide activity patterns.

Crucially, this approach allowed us to probe the relationship between the maze-position-specific coding subspaces defined by the set of SVM decoding weights associated with the activity of each pixel, for each task variable. We first plotted the pixel-wise weights across mice on a co-registered cortical space, with sub-region resolution. Interestingly, these weights were spatially heterogeneous across the dorsal cortex, often within single anatomical regions. They also varied with both task epoch and task variable, indicating dynamic contributions of different cortical (sub-)regions to each process (**Fig. 3d, Extended Data Fig. 5b**). To quantify how similar these coding subspaces were, we computed the angles between the decoding vectors defined by the pixel-level weights. In this analysis, angles close to 0° indicate highly similar coding spaces, while angles close to 90° indicate orthogonal, or linearly separable, subspaces. We used permutation tests to determine if these angles differed significantly from 90° (orthogonality) (**Fig. 3e**, absence of a dot indicates an orthogonal angle, see Methods for details). As with the PCA trajectories, the angles between coding subspaces for single task variables were significantly smaller within than across epochs (**Fig. 3e, f, Extended Data Fig. 5c, d;** results held when we used regression-based decoders for continuous behavioral variables, **Extended Data Fig. 6**). In other words, they were relatively stable within an epoch, but were mostly orthogonal across epochs. For instance, for ‘predominant color’ decoders this indicates that information about sensory evidence is stored in short-term memory in a separate subspace than the one used while the evidence was accumulated, which could protect this memory trace from interference of new (irrelevant) sensory stimuli (e.g., the wallpaper in the delay region). This is reminiscent of previously reported rotations of auditory information into an orthogonal subspace for memory storage within the auditory cortex^40^, and of a change in neural subspace in frontostriatal circuits when rats commit to a choice after accumulating evidence^41^. Finally, behavioral variables could also be decoded from pixels taken separately from within single cortical regions (**Extended Data Fig. 7**), consistent with the spatial heterogeneity of decoding weights across the dorsal cortex (**Fig 3d**). We also observed similarly orthogonal arrangements across epochs in these single-area decoders (**Supplementary Figs. 1, 2**), compatible with previous findings^14,21,40,42–52^ and suggesting that orthogonality is a scale-free property of cortical dynamics.

### Working-memory processes follow a spontaneous-timescale hierarchy

Our analyses so far show that the task’s decision variables are encoded by distributed, spatiotemporally dynamic activity patterns across the cortex. We next asked if we could make further sense of this large-scale arrangement. In particular, we hypothesized that the temporal evolution of the primary decision variable required to solve the task—sample category—would map onto a known principle of large-scale cortical organization, a hierarchy of spontaneous-activity timescales^53^. These timescales increase systematically from posterior sensory to frontal association regions and have been proposed to impact a region’s participation in cognitive computation^22,54–60^. For example, longer-timescale regions would be better suited to retain sensory information in memory in the absence of new input.

To test this, we first estimated the time course of sample-category information separately for each cortical area, by training separate ‘predominant color’ SVM decoders using pixel-level activity from within each anatomically defined region. Confirming the distributed nature of the representation, all single-area decoders performed above chance, albeit with varying accuracy (**Fig. 4a**). We then separately projected area activity onto its decoding axis for each behavioral trial (**Fig. 4b**). This has been interpreted as a single-trial read-out of (one-dimensional) decision variables^61–63^. For most trials, we found that the magnitude of projected activity, i.e. the amount of information about sample category, tended to increase over the sample region, and decrease after delay onset. We quantified this time course by separately fitting lines to single-trial projection magnitudes within the sample or delay regions (**Fig. 4b**, lines outperformed exponentials in a preliminary cross-validated model comparison). Interestingly, regions differed significantly in terms of the rate of change in their decision variables (i.e., the slopes of the linear fits, **Fig. 4c–e**, *P ≤* 0.001, one-way repeated-measures ANOVA; for other task variables, see **Extended Data Fig. 7, Supplementary Fig. 3**). Specifically, the decision variable rose fastest during evidence accumulation in posterior regions and decayed slowest in frontal regions during the delay. This was not due to stronger responses to high luminance in visual areas, since this was also present in trials with predominantly black checkerboards (**Fig. 4c–e**) and regressing out luminance-related activity before the analysis yielded the same results (data not shown).

**Fig. 4.**
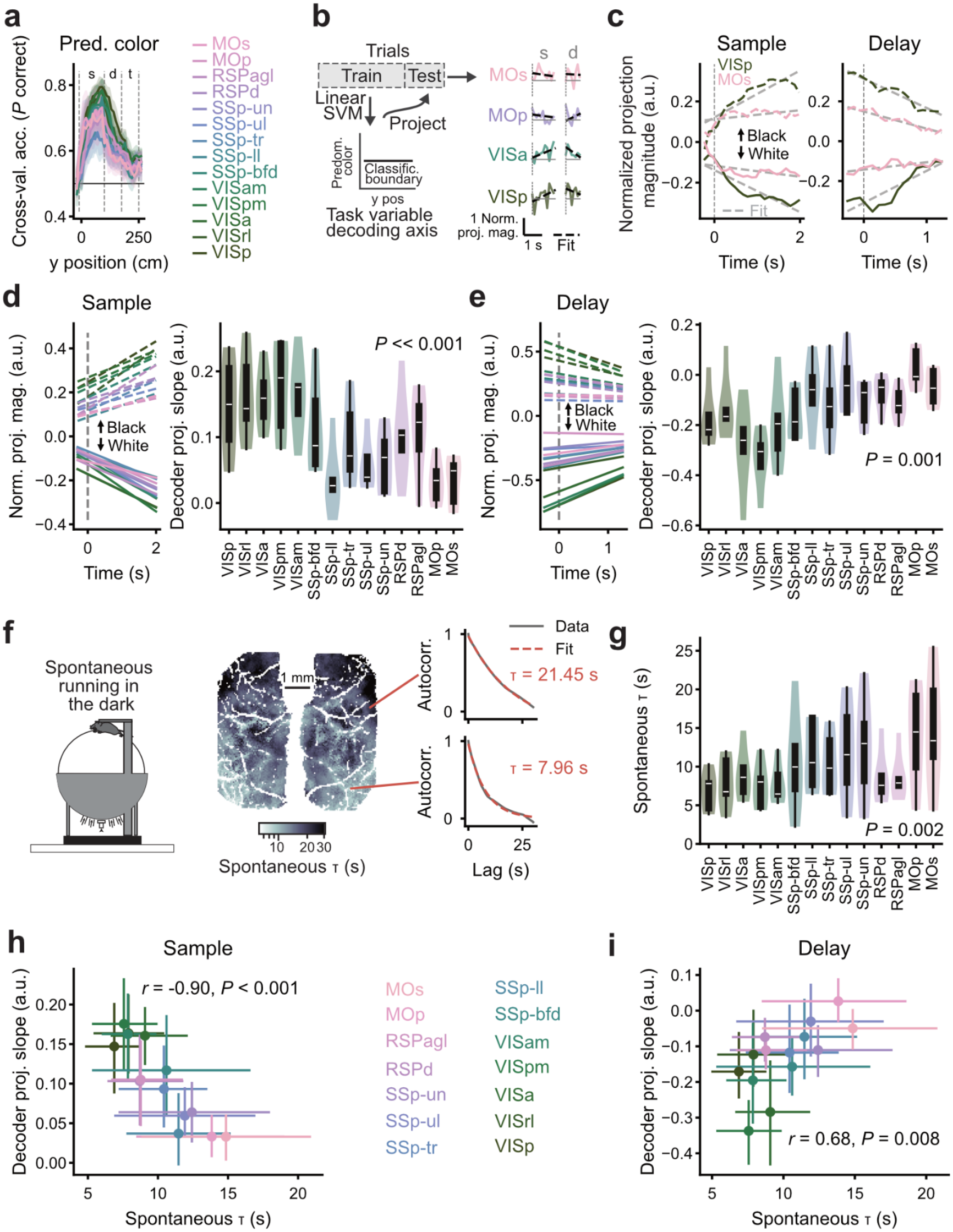
A hierarchy of spontaneous activity timescales predicts the time course of evidence accumulation and short-term memory in different cortical regions. **a**, Accuracy of decoders of predominant color from pixels taken separately from each cortical region, assessed on held-out trials. Different decoders were trained for each session, cortical region and y position (*n* = 20 sessions, 5 mice). 67–315 trials per session were split into training sets (*n* = 53–252 trials per set) and test sets (*n* = 14–63 trials per set). Error shades: 95% CI across sessions (*n* = 20). s: sample, d: delay, t: test. **b**, Schematics of projection-slope analysis. For each held-out trial, we projected the activity of pixels within an area onto the single-area decoding axes, obtaining single-trial estimates of decision variables. We then fit a line to the projected activity, separately for each trial and sample and delay epochs. **c**, Average projections during the sample (left) or delay (right) epochs for decoders trained on pixels from two example areas, shown separately for trials with predominantly black (dashed lines) or white (solid lines) sample stimuli. Dashed gray line: linear fit. **d**, Average fits for each area across held-out trials during the sample epoch (left), and unsigned fit slope distributions for each cortical region (right). Averages and violin plots are across subjects (*N* = 5). Printed *P*-value: one-way repeated-measures ANOVA. **e**, Same as panel d, for fits during the delay. **f**, Left: example pixel-wise cortical map of activity timescales measured while the same mice imaged during the task ran spontaneously in the dark, before task training. Right: example of autocorrelation functions and exponential fits used to quantify timescales (see Methods and **Extended Data Fig. 8** for details). **g**, Distribution of spontaneous timescales per anatomically defined cortical region. Printed *P*-value is from a one-way repeated-measures ANOVA, *n* = 5 sessions from 5 mice. **h**, Scatter plot of average area timescale vs. the average slope of projected activity during the sample. For each area, slopes from 755 held-out trials (14–63 per session) across 20 sessions (3–6 sessions per mouse) were averaged by mouse (*N* = 5) as in panels d and e above. Error bars: 95% CI across 5 mice. **i**, Same as panel h, for slopes during the delay.

To probe if the decision-variable time courses were indeed related to the timescale hierarchy, we measured mesoscale Ca^2+^ dynamics in a subset of the same mice in separate sessions in which they ran spontaneously in the dark (*n* = 5 sessions from 5 mice). Crucially, these sessions happened before task training. To estimate the spontaneous timescale of each pixel, we first regressed out running-related neural activity using ridge regression, and computed the autocorrelation function of the residuals of that model. We then performed model selection between a single exponential decay function or the sum of two exponentials, extracting the decay time constant from the best model (*τ*). Simulations confirmed that, compared to simply fitting a single exponential to an autocorrelation function of the full signal, this strategy more accurately recovered ground-truth timescales and was more robust to different levels of noise (though it slightly overestimated timescales due to GCaMP dynamics; **Extended Data Fig. 8**). Compatible with previous reports^22,55–57^, we observed increases in spontaneous timescales going from posterior sensory to frontal regions (**Fig. 4d, e**, *P* = 0.002, one-way repeated-measures ANOVA). Interestingly, timescale values were much larger than those previously reported for single neurons, even measured with cellular-resolution Ca^2+^ imaging (e.g., ^55,64,65^). This likely reflects our recent finding that single neurons combine into much longer-lasting population dynamics in the cortex^65^.

Next, to examine how the time course of the decision variable within a region related to its spontaneous timescale, we plotted the average timescale of each dorsal cortical region against the slope of activity along the ‘predominant color’ coding subspace. We found a strong negative correlation for the sample epoch (**Fig. 4f**, *r* = -0.90, *P* < 0.001, linear correlation coefficient), which suggests that short-timescale areas accumulate sensory evidence faster than long-timescale areas, similar to previous conclusions in mice and humans^22,66^. On the other hand, another recent study in mice found that, while cortical regions differ in their latency and duration of responses to evidence pulses, these differences do not strictly follow the spontaneous hierarchy^9^. However, differences in the stimuli (gratings vs. checkerboard vs. evidence pulses), methods of timescale estimation, and task design between the different studies complicate direct comparison. Moreover, cognitive processes themselves have been shown to modulate activity timescales^67^. Spontaneous timescales aside, all studies observed increasing evidence-integration timescales over multiple stages of cortical processing, potentially compatible with theoretical models in which integration is implemented via successive steps of feedforward processing^15,68^.

Finally, opposite to the evidence epoch, delay-epoch slopes were positively correlated with spontaneous timescale (**Fig. 4g**, *r* = 0.68, *P* = 0.001). Coupled with primarily negative slope values, this finding suggests that long-timescale areas retain accrued evidence for longer, compatible with previous evidence from the monkey cortex^10^. Importantly, these effects were largely specific to the predominant-color (and coherence) decision variable (**Extended Data Fig. 9**). Altogether, our findings show that spontaneous-timescale hierarchies are specifically predictive of both sensory evidence accumulation and short-term memory maintenance. This suggests that working-memory dynamics could be partly determined by intrinsic microcircuit properties and interareal connectivity^69–71^.

### Coding subspaces for different cognitive processes are orthogonal to each other

Our decoding analyses show that individual cognitive processes unfold within time-varying coding subspaces across the cortex. This highly dynamic arrangement for many concurrent cognitive variables could lead to their entanglement, i.e., at any given point in time more than one process could occupy a similar subspace to another. On the other hand, recent work suggests that, within single regions, different behavioral processes correspond to orthogonal activity patterns, which could minimize interference between them^14,21,40,42–52^.

To answer how the task’s different processes are arranged relative to each other at large scales, we leveraged the cortex-wide decoders from previous analyses (**Fig. 3, Extended Data Fig. 5**). We again computed the pairwise angles between the maze-position-dependent decoding axes defined by pixel-level SVM weights, but this time between different task variables. Notably, we found that all variables were fully orthogonal throughout the trial (**Fig. 5**). This is consistent with our previous visualization of the cortical maps of decoder weights, which appeared markedly distinct between task variables (**Fig. 3d, Extended Data Fig. 5b**). Thus, although the subspaces for a given variable changed across the trial, they remained orthogonal to subspaces spanned by other task variables. In other words, the cognitive processes in the task are disentangled at cortex-wide scales.

**Fig. 5.**
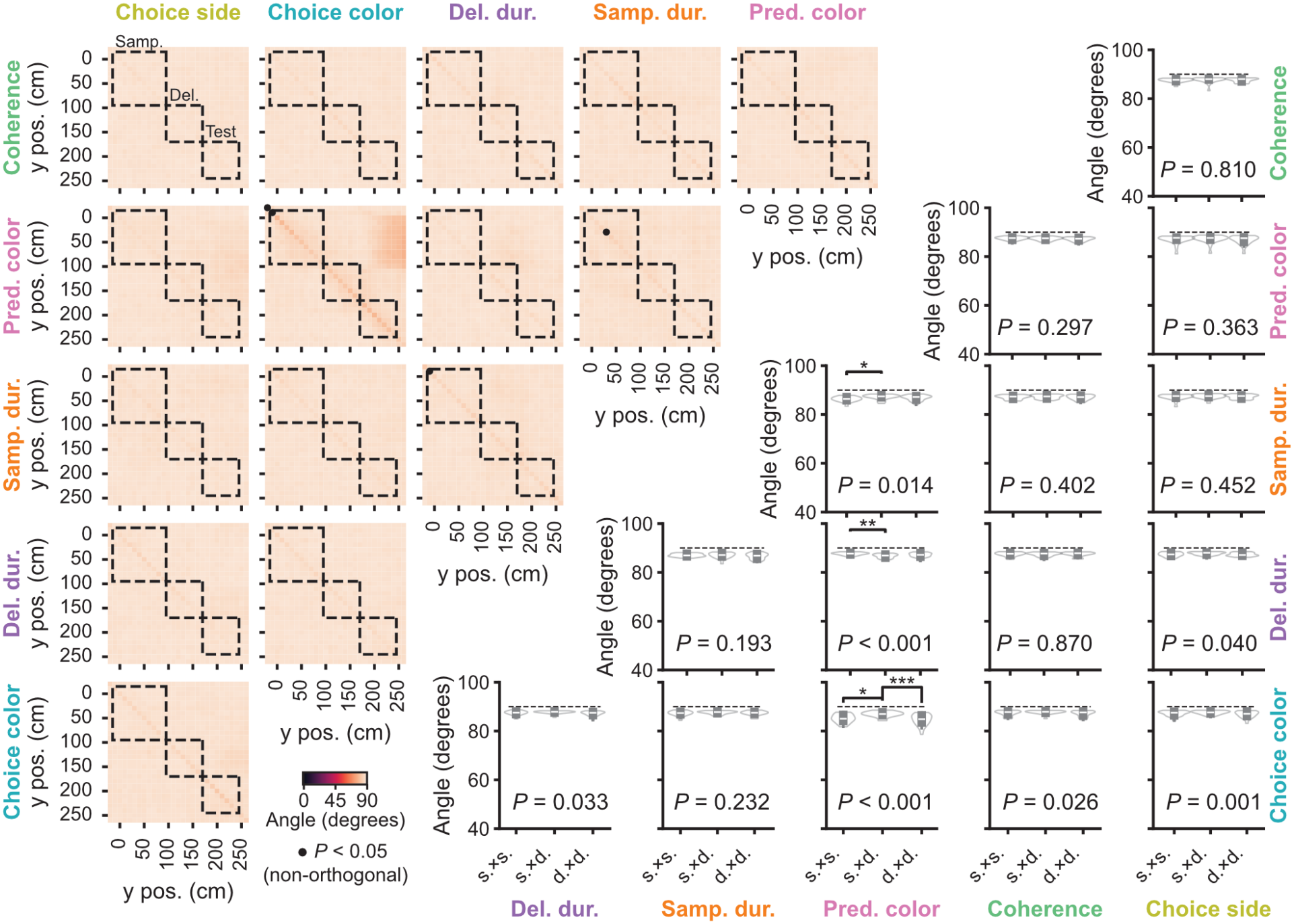
Coding subspaces for all task variables are orthogonal to each other. Top left: average angles between decoding axes for pairs of task variables by maze position. Dashed boxes: task epochs. Circles: significantly non-orthogonal angles (*P* < 0.05, shuffle test, see Methods for details). Angles and *P*-values are averages across 20 sessions from 5 mice. 67–315 trials per session were split into training sets (*n* = 53–252 trials per set) and test sets (*n* = 14–63 trials per set). Note that nearly all position-wise comparisons across task variables are orthogonal. Bottom right: decoding-angle distributions across sessions for task-epoch comparisons between pairs of task variables (*n* = 20 sessions). Printed *P*-value is for a one-way repeated-measures ANOVA with factor ‘epoch pairs’. *: *P* < 0.05, **: *P* < 0.01, ***: *P* < 0.001, Tukey’s post-hoc test. Note only a few significant differences between within- and across-epoch comparisons.

### Large-scale geometry is disrupted in error trials

If the geometric arrangement we uncovered is functionally relevant, it should be related to task execution. Specifically, if orthogonality prevents interference between cognitive processes, non-orthogonal arrangements might be associated with poorer performance. To test this, we first trained cortex-wide SVM decoders separately on correct and error trials (see Methods for details on trial-count matching), hypothesizing that the angles between the decoders for different task variables would decrease significantly during errors. This is in fact what we observed not only between variables, but also for between different maze positions within a variable (**Fig. 6a–c, Extended Data Fig. 10a**). In other words, in error trials, cognitive processing was both less dynamic and less distinct between different behavioral variables.

**Fig. 6.**
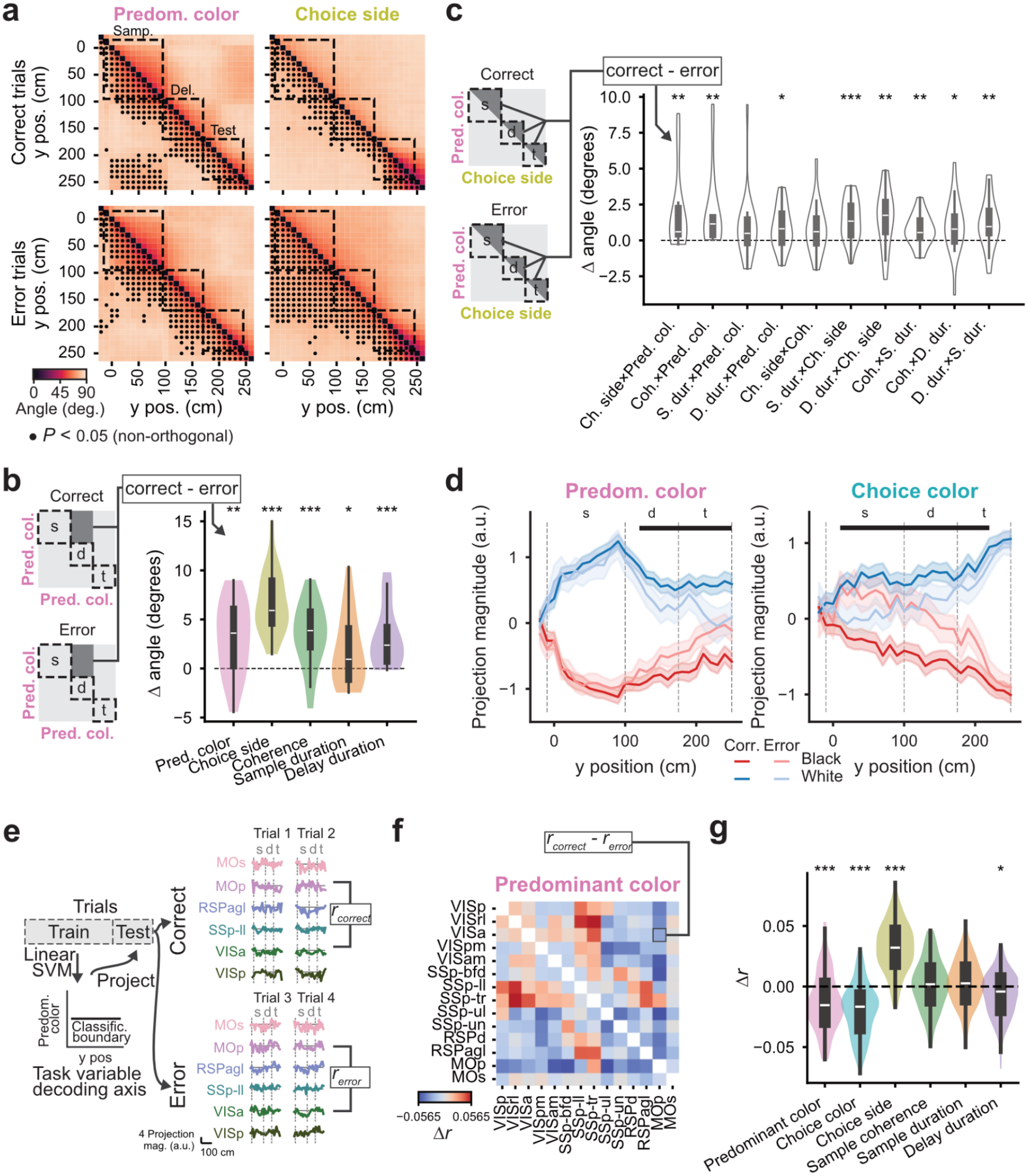
Coding subspaces are dynamic and break down during error trials. **a**, Angles between the decoding axes for the same behavioral variable at different maze positions for decoders trained separately on correct and error trials. Dashed boxes: task epochs. Circles: significantly non-orthogonal angles (*P* < 0.05, shuffle test, see Methods for details). Angles and *P* -values are averages across 20 sessions. Correct (*n* = 2539, 40–248 trials per session) and error (*n* = 1192, 21–109 trials per session) trials were count-matched to exactly 300 trials per session, then split into training (80%, *n* = 240 trials per session) and test (20%, *n* = 60 trials per session) sets. See Methods for details on trial-count matching. **b**, Difference between the angles of decoders trained separately on correct or error trials, across the sample and delay epochs for each task variable (*n* = 20 sessions). *: *P* < 0.05, **: *P* < 0.01, ***: *P* < 0.001, paired *t-*test between correct- and error-trained decoder angle averages across the shaded region in the schematic, with FDR correction (*n* = 20 sessions). **c**, Same as panel b, but comparing within epochs across different pairs of behavioral task variables. **d**, Cortex-wide activity projected onto the ‘predominant color’ (top) or ‘choice color’ (bottom) decoding axes, separately for correct and error trials (*n* = 494 correct, 261 error held-out trials, combined across sessions). Error shades: 95% CI over trials. Horizontal bars on top: *P* < 0.05, permutation test (correct vs. error trials, combined for black and white predominant or chosen color). **e**, Schematics of projection-correlation analysis. For each trial, we projected the activity of pixels within an area onto the single-area decoding axes, obtaining single-trial estimates of decision variables. We then computed the pairwise linear correlations between decision variables over maze position, separately for correct and error trials. **f**, Trial-averaged difference in the pairwise area correlations between correct and error trials (Δr) for ‘predominant color’ decision variables (*n* = 494 correct, 261 error held-out trials, combined across sessions). **g**, Distribution of Δr for different behavioral variables over area pairs (*n* = 91). For visualization purposes only, we first averaged pairwise area correlations between correct and error trials within each session (*n* = 20), subtracted the averages, and then averaged these differences across sessions, yielding a single datapoint for each area pair. For statistical analysis, we used linear mixed-effects models comparing session-averaged pairwise area correlations between correct and error trials for each task variable, with random effects for area pairs (*n* = 91), session (*n* = 20) and mouse (*N* = 5). ***: *P* < 0.001, *: *P* < 0.05, from the mixed-effects model.

Next, to analyze how this more entangled arrangement in error trials impacted the encoding of information required to solve the task, we projected cortex-wide activity onto the decoding axes for ‘predominant color’ separately for correct and error trials (using decoders trained on both trial types). This allowed us to examine the time course of activity specifically within the subspace spanned by this task variable (similarly to **Fig. 4**). In correct trials, activity encoding white or black predominant colors remained well separated throughout the maze. Conversely, in error trials, although this separation was preserved during the presentation of the checkerboard, it decayed throughout the delay and test stimulus epochs, suggesting that checkerboard category was encoded normally but then forgotten (**Fig. 6d**, left). Similar conclusions can be drawn from projecting activity on the ‘choice color’ decoding axis—activity initially separated black and white with signs compatible with a correct upcoming choice, but this eventually decayed and reversed to the erroneous choice by the delay region (**Fig. 6d**, right). Interestingly, these differences in projection magnitude between correct and error trials were specific for these two behavioral variables (**Extended Data Fig. 10b**). Moreover, these effects were not trivially related to error trials having longer delays on average, since they were true regardless of sample and delay duration (**Extended Data Fig. 10c**).

Lastly, we wondered whether errors could also be related to changes in the relationship between cognitive processing in different regions. Specifically, because decreases in large-scale correlations have been associated with increased cognitive demands^7,23,72,73^ and improved task-information coding^74^, we hypothesized that error trials would be associated with higher correlations between regions, within the subspaces used for cognitive computation. To get at this, we used the same single-area decoders from **Fig. 4** and separately projected single-area activity onto its decoding axis to obtain decision variables (**Fig. 6e**). We then computed the correlation between decision-variable time courses across pairs of areas and we compared these correlations between correct and error trials. Correlations were significantly larger in error trials (**Fig. 6f, g, Extended Data Fig. 10d**), specifically for ‘predominant color’ (*P* < 0.001, linear mixed-effects model), ‘choice color’ (*P* < 0.001), and ‘delay duration’ (*P* < 0.05). The interesting exception was ‘choice side’, which was significantly more correlated in correct trials (*P* < 0.001), compatible with a recent report on the human cortex^61^. Altogether, our findings suggest that error trials are associated with more entangled cognitive processing at cortex-wide scales, supporting the functional significance of the large-scale orthogonal arrangement that we have uncovered.

## Discussion

Here, we developed the first decision-making task for mice that dissociates evidence accumulation, sensory short-term memory, and motor choice (**Fig. 1**). This task allowed us to characterize how cortical dynamics relate to each cognitive process, and how process-specific dynamics co-exist at cortex-wide scales. We found that, although each cortical area contains information about every cognitive and non-cognitive component of the task (**Fig. 2**), each cognitive process occupies dynamically changing orthogonal subspaces defined by multi-area activity (**Figs. 3, 5**). Crucially, the coding subspaces for sensory evidence accumulation and short-term memory appear to co-opt the cortex’s intrinsic organization along a gradient of spontaneous timescales (**Fig. 4**). Finally, we showed that coding subspaces are less dynamic and less orthogonal in error trials, and with increased correlations between single-area decision variables (**Fig. 6**). Thus, our work reconciles seemingly disparate findings on large-scale and single-area cortical dynamics by showing that locally orthogonal geometries are preserved into non-interfering global activity subspaces related to different cognitive processes.

At a minimum, the orthogonality of dynamics across task epochs and between task variables serves as a succinct, quantitative description of the finding that distinct activity patterns underlie different types of working memory—whether gradually updated via accrual or maintained over time to be used for an upcoming decision. The representational-geometry framework also proposes a mechanistic interpretation for it: a downstream region (neuron) receiving input from the population of neurons in question would be able to (linearly) decode these orthogonal patterns for behavioral purposes^42,45,46,48,50^. A similar logic could in principle apply to the present case, since several brain regions are known to receive converging projections from across the dorsal cortex, e.g., striatum^75–77^, medial prefrontal cortex^76,78,79^, superior colliculus^76,80,81^, all of which have been heavily implicated in working memory and decision making^6,9,82–89^. Future work should determine how this cortex-wide information is eventually combined towards a single behavioral output.

Upcoming studies should also probe the neurobiological mechanisms that generate globally orthogonal dynamics. Non-mutually exclusive possibilities include cortico-cortical connectivity and gradients of local cellular and circuit properties across the cortex^53,65,69–71^, cortico-thalamic and cortico-basal ganglia loops^82,90–93^, and neuromodulatory input to the cortex^73,94,95^.

Finally, beyond the conceptual advances above, we expect that our new task will spur work in the field at large, allowing new understanding about how different processes can be flexibly combined to enable complex cognitive behavior.

## Methods

### Animals and Surgery

All experimental procedures were approved by Northwestern University’s Institutional Animal Care and Use Committee and were done in accordance with the Guide for the Care and Use of Laboratory Animals. We used 10 mice of both sexes, aged 2–14 months (*N* = 7 for imaging experiments, and an additional 3 for just behavior). The imaged mice were obtained by crossing a flex-GCaMP6s line [Ai96, B6J.Cg-*Gt(ROSA)26Sor*^*tm96(CAG-GCaMP6s)Hze*^/MwarJ, JAX stock #028866] with a Cux2-Cre one [B6(Cg)-Cux2tm2.1(cre)Mull/Mmmh, RRID:MMRRC_032778-MU] to drive GCaMP6s expression preferentially in supragranular cortical layers. The three behavior-only mice were obtained by crossing three different lines with the flex-GCaMP6s line (Ntsr1-Cre [B6.FVB(Cg)-Tg(Ntsr1-cre)GN220Gsat/Mmucd, RRID:MMRRC_030648-UCD], PV-Cre [B6.129P2-*Pvalb*^*tm1(cre)Arbr*^/J, JAX stock #017320], and SST-Cre [STOCK *Sst*^*tm2*.*1(cre)Zjh*^/J, JAX stock #013044]). With very few exceptions, the mice were group housed and all were kept under a 12/12 h reversed light-dark cycle.

Each mouse underwent a single surgical procedure to implant a titanium headplate and optically clear the intact skull as described in detail previously^7^. Briefly, mice were anesthetized with isoflurane (3% induction, 1.5% maintenance) and positioned in a stereotaxic frame (Kopf). After asepsis, trichotomy and local application of lidocaine S.C. (4 mg/kg), the skin over the dorsal skull was removed and the periosteum scraped. Thin layers of cyanoacrylate glue (Elmers) and clear metabond (Parkell) were then successively applied. After curing, the metabond surface was polished with a cement polishing kit (Pearson Dental). The headplate was then bonded to the skull using opaque metabond. Finally, an even thin layer of transparent nail polish (Electron Microscopy Science) was applied over the implant. For analgesia, the mice received two doses of meloxicam (20 mg/kg S.C., perioperative and 24 h later). They also received warm saline after the procedure to maintain hydration (0.1–0.3 mL S.C). Body temperature was maintained at 37 °C throughout the procedure using a homeothermic blanket (Harvard Apparatus).

The mice recovered from the procedure above for at least 5 days before undergoing habituation to experiments. They were then water restricted to motivate behavioral training. Each mouse received a daily water allotment of 1–1.5 mL depending on their baseline *ad libitum* drinking weight, and supplemental fluids were provided if the weight fell below 80% of baseline. During the first ∼7 days of water restriction, mice were extensively handled and hand-fed their water allotment. Behavioral training started once the mice voluntarily climbed onto the experimenters’ hands and showed no signs of distress. After each behavioral training session, the mice had access to a large enclosure with toys and conspecifics for environmental enrichment.

### Behavioral task

#### Virtual-reality (VR) apparatus

The custom-built apparatus was similar to the one described in detail elsewhere^28^, modified for projection onto a more compact spherical screen. Head-fixed mice ran on a Styrofoam© ball suspended by compressed air (∼55 psi), and surface displacements on the ball were read out from a single optical velocity sensor lying underneath it (ADNS-3080 APM2.6), using an Arduino Due. These displacements were transformed into x-y and angular movements in the virtual world, in closed loop, using custom code written using ViRMEn, a MATLAB VR Engine^96^, running on a PC. View angle was computed as atan2(−d*X* × sign(d*Y*), |d*Y*|), where d*X* and d*Y* are the x and y displacements. The virtual world was projected onto a spherical screen (17” internal diameter, covering ∼200° horizontal and ∼70° altitude of the mice’s field of view) using a DLP projector (Optoma HD28HDR, 120-Hz refresh rate, image resolution of 768 × 1024 pixels, RGB color balance of [0 0.4 0.5]). Reward was delivered through a metal spout by opening a solenoid valve (NResearch) connected to a custom MOSFET circuit and controlled by a TTL from a DAQ card (National Instruments). The apparatus was enclosed in a custom-designed cabinet (The Knotts Company) fitted with sound-attenuating foam (McMaster Carr).

#### Task design

The task happened in a virtual T-maze with the stem measuring 10 × 300 × 5 cm and side arms measuring 20 × 20 × 5 (width × length × height). By convention, width corresponds to the x dimension of the sensor, and length to y. The length was divided into the following regions: start box (30 cm), sample region (100 cm), delay region (75 cm), test region (75 cm), and arm (20 cm). For two mice, the delay region was 50 cm, and the rest of the maze was identical. The transition zones between each region were indicated by distal landmarks hovering above the maze. At the beginning of each trial, the mouse was teleported to the start box and as it entered the sample region, a bilaterally symmetrical checkerboard pattern of black and white squares in different proportions was partially revealed on maze walls and floor (y = 20 cm, 10 behind and 10 in front of the mouse; the size of each square was ∼9 × 9 degrees of visual angle). This stimulus was revealed when the mouse was 10 cm away from it (and thus first visible at y = -10 cm), and the sliver of checkerboard moved with the mouse. After the delay region, two test stimuli were revealed, each on a side wall, one white and one black (to preserve optical-flow cues, these were implemented as a checkerboard with 95% of squares in the intended color). The mice were rewarded with a drop of 10% v/v sweet-condensed milk (4–8 µL) for turning towards the arm on the side matching the predominant color of the checkerboard. Errors were indicated with a chime played through speakers. Both a correct and a wrong choice resulted in the maze freezing for 1 s. The screen then went black for an inter-trial interval of 2 s or 10 s, respectively after a correct choice and an error. To enforce a symmetrical view of the checkerboard, the mouse’s virtual view angle was constrained to be zero until it reached the test portion of the maze, at which point view angle was released to allow turning. The exceptions were the two mice that ran on the 50-cm version, who did not have constrained view angles.

Three parameters were varied independently on a trial-by-trial basis. First, coherence of the sample, defined as the proportion of squares of the predominant color, was varied between 0.55 and 0.95, drawn from an exponential distribution with a rate parameter of -9, such that higher (easier) coherences were drawn more often than lower ones. Each square was independently generated by drawing from a binomial distribution with *P* equal to the generative coherence. The other two parameters were the gains applied to map physical to virtual x-y displacements, in order to vary the duration of the sample and/or delay regions (normal gain is 1). Gains were drawn from 3 discrete values, 0.67, 1, or 1.5, with fixed probabilities of 0.3, 0.4, and 0.3, respectively. Whether the sample was predominantly white or black, and whether the rewarded side was left or right, were both drawn from a dynamically varying *P* designed to counteract biases measured over the session (e.g., a left bias will result in an increased probability of drawing a right-rewarded trial, a white bias will similarly increase the probability of black-rewarded trials). The algorithm used to calculate *P* is described in detail elsewhere^28^. Finally, to maintain task engagement and motivation, the mice were occasionally exposed to a block of 10 easy trials, where the rewarded arm was indicated by a tall visual guide (70 cm) visible throughout the maze. This happened whenever performance over a 50-trial window (or 30, for sessions run in an earlier task version) fell below 55% correct. A typical behavioral session consisted of 200–300 trials, with a median trial duration of 7.3 s (trials longer than 60 s were automatically terminated and excluded from analysis).

#### Training procedure

After extensive habituation (see *Animals and Surgery*), the animals underwent initial motor training to control the treadmill by running on a 50-cm-long linear track. In this pre-training phase, mice were rewarded when they reached the end of the track in less than 60 s. Mice remained in pre-training until they could reliably obtain ∼1 mL reward in 20–30 min, which took a median of 4 sessions. They then entered a fully software-automated shaping pipeline of 13 steps, in which different aspects of the task were sequentially introduced once the mice reached a set of performance criteria. Details about the steps and criteria are provided in **Extended Data Fig. 1** and **Supplementary Tables 1–4**. Briefly, the first step consisted of a short maze with a persistent, 0.95-coherence sample checkerboard and test stimuli, a very short delay (10 cm), and a tall visual guide indicating reward location. The maze was lengthened over the first four steps, followed by removal of the visual guide to explicitly introduce the match rule (implicitly in place since step 1). The short-term memory requirement was then introduced by hiding the test stimulus until the end of the delay region, which progressively grew over the next few steps. Next followed the introduction of variable checkerboard coherence. Then, to enforce gradual evidence accumulation, the until-then fully visible checkerboard started being partially revealed as the mice ran. Finally, the last two steps consisted of the introduction of variable sample and delay durations from the gain manipulations, in this order.

### Mesoscale widefield Ca^2+^ imaging

Our custom-built apparatus for mesoscale-resolution widefield imaging consisted of a tandem-lens design (1× and 0.63× going from the sample to the sensor, Leica, M Series) with an intervening filter box (BrainVision Inc.) containing a dichroic beamsplitter (FF495-DI03-50X70, Semrock) and a green emission filter (FF01-525/45, Semrock). Images were captured at 20 Hz and a resolution of 1024 × 1024 pixels (pixel size of 6.8 µm, field of view: ∼7 × 7 mm) using a Prime BSI Express sCMOS (Teledyne) and the open-source software µManager running on a PC. To allow for hemodynamic contamination correction using the isosbestic wavelength, illumination alternated between a blue and a violet LED (Cairn, 470 and 405 nm, respectively, each bandpass-filtered at a width of 40 nm and driven by a MultiStream Pro, power of ∼2 mW/cm^2^ at the sample). The alternation was controlled by exposure signals from the sCMOS via custom code running on an Arduino Due. Illumination was delivered to the filter cube using a liquid light guide (½” diameter, Edmond Optics). The objective was light-shielded from the VR projection using a custom 3D printed part. Imaging signals were synchronized with behavior by directly reading exposure signals from the sCMOS using a DAQ card (National Instruments) on the VR PC, and saved directly into the behavioral data file.

### Histology, immunohistochemistry, and ex-vivo imaging

We verified the laminar distribution and cell-type specificity GCaMP expression with immunostaining in a subset of mice. They were deeply anesthetized with ketamine (50-100 mg/kg) and xylazine (10 mg/kg) and transcardially perfused with phosphate-buffered saline (PBS, 20 mL), followed by 4% paraformaldehyde (10 mL) for tissue fixation. The brains were then removed and post-fixed in 4% w/v paraformaldehyde for 12–24 h, followed by cryoprotection with 30% w/v sucrose. The tissue was coronally sliced in 30-µm sections using a cryostat (Leica) and slices were transferred into wells for staining. The slices were washed with PBST (0.4% Triton-X in PBS, 3 × 10 min), incubated with blocking buffer for 2h at room temperature (2% v/v normal donkey serum + 10 mg/mL bovine serum albumin), and incubated overnight with primary antibody at 4 °C. This was followed by another PBST wash (3 × 10 min), 2 h of incubation with secondary antibody at room temperature, a final PBST wash (3 × 10 min), and mounting on glass slides with Fluoromount-G™ with DAPI. In most brains, we stained for GFP only—primary antibody: anti-GFP antibody chicken polyclonal (1:4000, Abcam, ab13970); secondary: Alexa Fluor 488 Donkey Anti-Chicken IgY (1:1000, Jackson ImmunoResearch, 703-545-155). In one brain, we additionally stained for CaMKIIα to verify that GCaMP expression was restricted to excitatory neurons. In that case, besides the antibodies above, we incubated the samples with the following—primary antibody: mouse anti-CaMKIIα (1:50, Santa Cruz Biotechnology, sc-13141); secondary: Alexa Fluor 647 Donkey anti-Mouse IgG (1:200, ThermoFisher, A-31571). Slices were imaged using a confocal microscope (Nikon).

### Behavioral analysis

*Data selection*. Because of the easy-trial blocks, behavioral sessions were naturally divided into variable-length trial blocks of the main task. We selected for analysis entire blocks of trials with block-wise performance ≥ 0.6 fraction correct trials. Individual trials were then excluded if they reached the maximum duration of 60 s or if the total distance traveled exceeded 110% of the nominal maze length. These criteria yielded 49800 trials from 454 sessions in 10 mice for behavioral analysis (**Fig. 1**). The same criteria were applied for the selection of neural data. For time-based analyses (**Fig. 2, Extended Data Fig. 3**), we used data from all mice (*n* = 5411 trials, 29 sessions and 7 mice), while for space-based analyses (**Figs. 3–6, Extended Data Figs. 4–7, 9, 10, Supplementary Figs. 1–3**) we excluded the mice that ran with shorter delay regions (yielding *n* = 3731 trials, 20 sessions and 5 mice).

#### Performance metrics

Because task parameters were defined on a continuous space, performance was quantified by binning coherence, sample duration, and delay duration into evenly spaced bins in either a linear (coherence) or logarithmic (sample and delay durations) scale, excluding bins with fewer than 20 trials. Within each bin, fraction correct was defined as the proportion of trials in which the mice turned to the side whose test color was the same as the predominant color of the sample. Side bias (used in shaping-step advancement criteria, **Supplementary Table 3**) was defined as the difference between fraction correct for right and left rewarded trials.

#### Behavioral generalized linear models (GLM)

The Bernoulli GLMs in **Fig. 1e, Extended Data Fig. 2d–g** were trained separately for each mouse using 10-fold cross-validation and L2 regularization. Reported accuracies are on held-out test trials. Correct and error trial numbers were matched by subsampling. In addition, for models predicting choice side or color (**Extended Data Fig. 2d, e**), the numbers of trials were balanced for the predicted variable using subsampling. All predictors were z scored. The choice-accuracy models were formulated as:

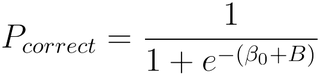

Where *β*_0_ is an offset term and was as follows for **Fig. 1**:

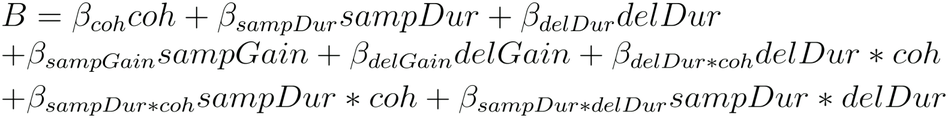

Where *coh* is the empirical checkerboard coherence for a given trial, *sampDur* is sample duration, *delDur* is delay duration, *sampGain* is the gain in the sample region, *delGain* is the movement gain in the delay region, and denotes interaction. Note that, because the gain is discrete but speed follows continuous and only partially overlapping distributions (**Fig. 1a**), the gain and duration terms are orthogonal, so the design matrix is still full rank. We included both terms in the model to account for the possibility that the gain itself affects performance independent of duration (e.g., by changing running effort). The model in **Extended Data Fig. 2f** was identical to the model above, with the addition of the running speed term +*β*_*speed*_*Speed* to the right-hand side of the equation.

For the model in **Extended Data Fig. 2g**, *B* was defined as:

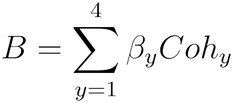

Where are equally spaced y position bins within the sample region of the maze and *Coh*_*y*_ is the empirical checkerboard coherence for that maze segment and trial.

Finally, the trial-history models in **Extended Data Fig. 2d, e** were parameterized as:

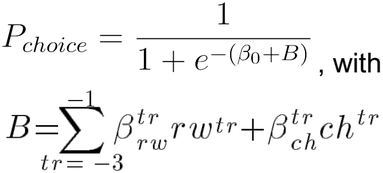

Where *rw*^*tr*^ and *ch*^*tr*^ are respectively the reward indicator function and the choice side (or color) preceding trial. In the choice-side model *P*_*choice*_ corresponds to a leftward choice, and to a white choice in the choice-color model.

### Widefield Ca^2+^ imaging preprocessing

Our preprocessing pipeline for widefield imaging data was a modified version of our published one^7^. Image stacks were first corrected for x-y motion by applying the translation that maximized the phase correlation between each frame and the average across frames. The frames were then spatially binned to 128 × 128 pixels (resulting pixel size of 54.4 µm) and parsed into blue- and violet-excitation frames. Separate Δ*F*/*F* traces for each excitation wavelength, Δ*F*/*F*_*B*_ and Δ*F*/*F*_*V*_, were then computed as (*F* − *F*_0_)/*F*_0_, where *F*_0_ was defined as the mode of *F* for Δ*F*/*F*_*B*_ and the 80^th^ percentile of *F* for Δ*F*/*F*_*V*_, both computed over a 30-s running window. We then obtained a scaled version of Δ*F*/*F*_*V*_, 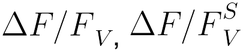, by using linear regression to find a single scalar slope and offset that optimally matched the amplitudes of the troughs in Δ*F*/*F*_*B*_ and Δ*F*/*F*_*V*_ (i.e., by thresholding the traces to only have values smaller than 0). This and the choice to model *F*_0_ as the 80^th^ percentile for violet excitation are both based on the intuition that hemodynamic contamination should primarily lead to decreases in fractional fluorescence^97^. Indeed, simulations confirmed that these parameters lead to better correction (not shown). Next, we smoothed 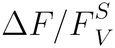 with a 400-ms Gaussian kernel. Finally, was computed as a divisive correction given the multiplicative nature of hemodynamic contamination^97^, i.e., 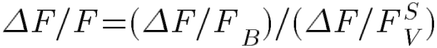. Because large blood vessels have different hemodynamic properties that make them harder to correct^7,97^, we masked them using the following algorithm. We computed the deviation of an unbinned average frame (i.e., 1024 × 1024 pixels) from a smoothed version of this average frame obtained by adaptive median filtering. We then masked any pixels whose deviation exceeded 1.6 in the negative direction or 10 in the positive direction. For sessions where mice ran spontaneously in the dark (see *Estimation of spontaneous timescales and related simulations* below), we combined this vasculature mask with a second mask computed using the same method, but in which the first principal component of the detrended raw fluorescence was used instead of the average frame. This automatically computed vascular mask was then combined with a manual mask applied to off-implant pixels (e.g., headplate). Masked pixels were assigned NaN values for all subsequent analyses.

#### Registration to brain atlas

To group data into anatomically defined cortical areas, we fitted the average frame of each session to a horizontal projection of the Allen Brain Atlas CCFv3, using a rigid transformation with scaling between two fiducial markers: lambda and bregma. The horizontal projection was obtained by selecting the most dorsal voxel from layer 2/3 for each 50 µm bin in atlas space.

#### Activity binning by maze position

To ensure equal data occupancy across bins for analyses in maze space (**Figs. 3–6, Extended Data Figs. 4–7, 9, 10, Supplementary Figs. 1– 3**), each 10-cm spatial bin consisted of the activity observed at the time the mouse first entered the bin (although mice rarely entered the same bin multiple times). We only considered data points within the trial (i.e., after teleportation and before trial-feedback delivery).

### Linear encoding models of neural activity

We fitted a linear mixed-effects encoding model to area-averaged data (**Fig. 2f, g, Extended Data Fig. 3, Supplementary Table 5**). For each anatomically defined unilateral area (*n* = 28), we averaged pixel *ΔF/F* in the native temporal resolution of the data (10 Hz). We then z-scored the data within sessions and concatenated trials from all sessions and animals to fit single models to each area. Prior to fitting, the concatenated response variable was z-scored a second time. We only considered time points between the start box and the test stimulus (i.e., we excluded the arm region and the inter-trial interval). The models were fitted with L2 regularization and 5-fold cross validation, with splits at the level of whole trials, not time points, to account for the temporal autocorrelations introduced by the Ca^2+^ sensor when estimating goodness of fit. Splits were also stratified to ensure proportional participation of each imaging session in cross validation. Trials were balanced for black and white choice color. After a preliminary round of cross-validated model comparisons, the final model was parameterized as follows:

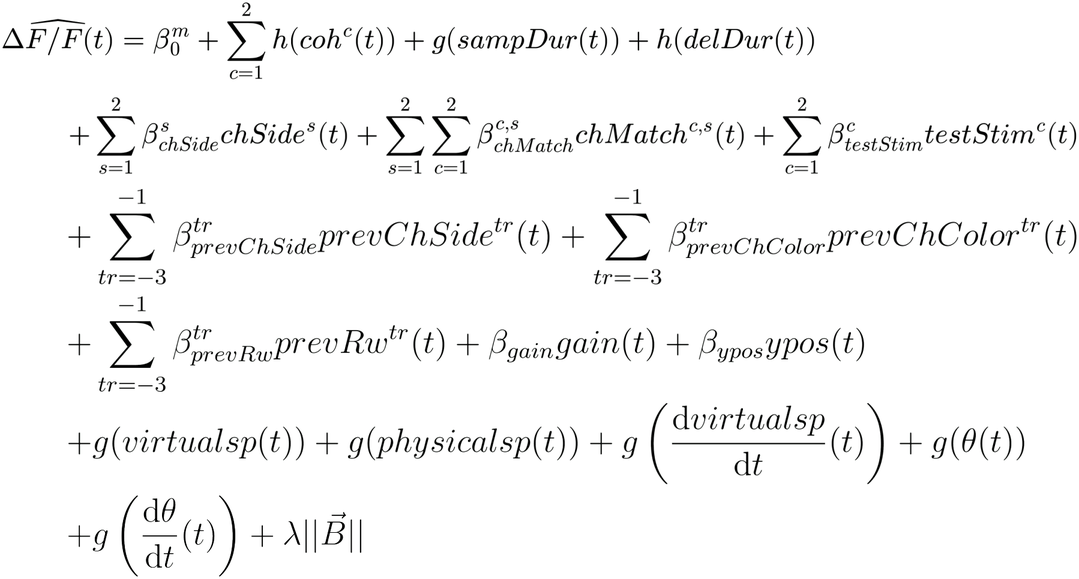

Where 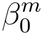, the mixed-effects term, is a random offset for each mouse, modeled as a multivariate Gaussian:

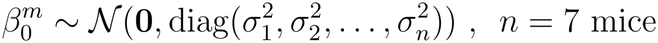

*g*(*x*) is a polynomial expansion of the predictor *x*(*t*) up to the 5^th^ degree, with degree 0 corresponding to a step function:

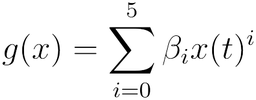

*h*(*x*)is a convolution of *x*(*t*) with a 3^rd^-degree basis set of 7 splines 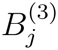 over 2 seconds:

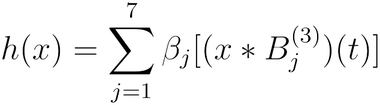

And *coh*^*c*^(*t*) are the sample coherences modeled separately for each color (black or white) during the sample region (and 0 otherwise); *sampDur(t*) is the cumulative elapsed time within the sample region (and 0 otherwise); *delDur(t*) is the same but for the delay region; *chSide*^*s*^(*t*) are indicator functions for upcoming choice side *s* (left or right), present throughout the trial; *chMatch*^*c,s*^(*t*) are indicator functions for match vs. no-match choice for the different color and side combinations (e.g., match black on left), during the test region (and 0 otherwise); *testStim*^*c*^(*t*) is the stimulus pattern *c* during the test region (e.g., for *c*=1 the right-side wall and floor are black and the left-side wall and floor are white, whereas for *c=2* the colors are flipped); *prevChSide*^*tr*^(*t*) and *prevChColor*^*tr*^(*t*) are single indicator functions for chosen sides and colors in the three previous trials *tr*, present throughout the current trial; *prevRw*^*trial*^(*t*) are analogously indicator functions for whether the animal was rewarded in the three previous trials; *gaint (t*) is the instantaneous virtual movement function gain; *virtualsp* is the linear x-y speed in the virtual environment; *physicalsp(t*) is the physical running speed extracted from the movement sensor (different than *virtualsp*(*t*) when *gaint*(*t*) ≠1); *dvirtualsp/dt(t*) is linear acceleration; θ(*t*) is the virtual view angle; and *dθ /dt*(*t*) is virtual angular velocity. Finally, 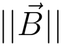 is the L2-norm of the coefficient vector and is the regularization parameter, determined by cross-validation.

For the reduced model analysis, we dropped entire groups of predictors and refitted the model. For instance, the model without delay duration dropped the entire spline basis set, and the model without kinematics excluded all speed and view angle predictors. We then quantified the contributions by subtracting the full model *R*^*2*^ from the reduced one. We computed the significance of the reduction by calculating an *F* statistic and extracting the *P*-value from an *F*-distribution with the appropriate degrees of freedom, on the held-out trial set:

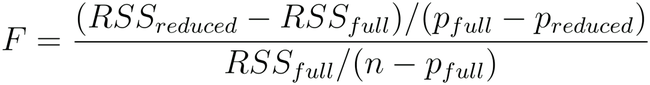

where *RSS*_*readuced*_ and *RSS*_*full*_ are the residual sum of squares, *p*_*readuced*_ and *p*_*full*_ are the number of parameters, and *n* is the number of observations (data points).

### Principal components analysis (PCA)

For the analyses in **Fig. 3a, b** and **Extended Data Fig. 4a–c**, for each selected trial within a behavioral session we computed activity binned by maze y position as described above. Trials within each session were then concatenated to form a matrix of dimensions *position bins*trials* × *pixels*. This matrix was z-scored and decomposed using the randomized singular value decomposition method^98^. The average trajectories in **Fig. 3a, Extended Data Fig. 4b** were obtained by projecting each trial onto the top 3 PCs and averaging across data points. Angles between pairs of N-dimensional trajectory segments were obtained by subtracting the start point of each segment from its end point, and then computing the arccosine of the dot product between the end points. The analyses in both **Fig. 3b** and **Extended Data Fig. 4c** were performed on the top 20 PCs, although selecting different numbers of PCs yielded qualitatively similar results.

#### Regressing out kinematics and luminance

The trajectories used in **Extended Data Fig. 4b, c** were computed on residual pixel-wise activity after regressing out contributions from position, speed, acceleration, view angle, view angle change, treadmill velocity, and luminance for the left and right hemifields. Luminance traces were calculated by reconstructing the visual scene in front of the mouse at each iteration of the VR engine (ViRMEn) for every trial. The visual scene was reconstructed from the logged behavioral data, the visual and geometric properties of the virtual maze, and the geometry of the spherical screen projection. The luminance of each hemifield was then computed as the sum of all pixels and channels on each side of the screen. The L2-regularized linear encoding model fit to the activity of each pixel from each session was:

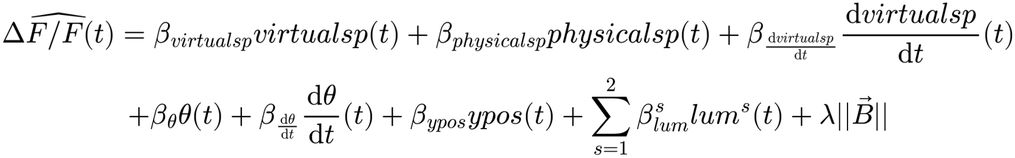

Where *virtualsp* is the linear x-y speed in the virtual environment; *physicalsp(t*) is the physical running speed extracted from the movement sensor (different than *virtualsp(t*) when *gaint(t) ≠*1);*dvirtualsp/dt(t*) is linear acceleration; *θ(t*) is the virtual view angle; *dθ/dt(t*) is virtual angular velocity; *ypos* is the y position in the virtual maze; and *lum*^*s*^ is the luminance of each hemifield *s* . Finally, 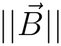 is the L2-norm of the coefficient vector and is the regularization parameter, determined by cross-validation.

### Support vector machine (SVM) decoding models

For the SVMs in **Fig. 3–6, Extended Data Figs. 5, 7, 9, 10**, and **Supplementary Figs. 1–3**, we created binary categories for the continuous variables using median splits as above. We then trained a soft-margin linear classifier SVM with L2 regularization and 5-fold cross-validation, separately for each session (regularization parameter = 1, soft margin = 0.1). For the regression SVM decoder in **Extended Data Fig. 6**, we binned continuous variables in five percentile bins, then trained a soft-margin linear regression SVM with L2 regularization and 5-fold cross-validation separately for each session (regularization parameter = 0.0001, soft margin = 0.1). Angles and cross-validated accuracy were averaged across sessions. All models were trained on images spatially binned to a resolution of 32 × 32 pixels. For weight plots (**Fig. 3d, Extended Data Figs. 5b, 6b**), these ∼200-µm binned pixel coefficients for each session were aligned back to the Allen Brain Atlas CCFv3 (as described in *Widefield Ca*^*2+*^ *imaging preprocessing*) with 50-µm resolution in atlas space. For visualization, we only plotted atlas bins appearing in at least 8 sessions. Note that, because the weight maps are first registered in high resolution before averaging across sessions, they may be offset relative to one another by less than the ∼200-µm binned resolution, leading to an appearance of finer resolution. In other words, this is an artifact of plotting only. The angles between the decoding axes were computed as the arccosine of the dot product between the weight vectors, and tested for orthogonality using the permutation test described below. For analyses involving projections onto the decoding axis (**Figs. 4b–e, h, i, 6d–g, Extended Data Fig. 9, 10a, c, d**), we only analyzed the projections of the 20% of held-out trials not used to fit the model, and combined trials across sessions for statistics. For the statistical comparisons between projections for correct and error trials (**Fig. 6d, Extended Data Fig. 10a, c**) we used a two-tailed permutation test to determine whether the difference we observed was greater than the null distribution obtained from computing the difference between 9999 random pairs resampled from the pooled correct and error time courses. Finally, we applied a threshold requiring at least two consecutive significant y positions using an algorithm adapted from^99^. To allow for a single comparison between correct and error trials, the sign for one of the variable’s categories (e.g., white sample color) was flipped, and both categories were combined. Trial-by-trial correlations between the area projections (**Fig. 6f, g, Extended Data Fig. 10d**) were computed separately for each area pair and then averaged across either trials for each session (**Fig. 6g**) or trials combined across all sessions (**Fig. 6f, Extended Data Fig. 10d**).

### Permutation tests for orthogonality

We followed procedures similar to previous publications to determine if the angles differed significantly from 90° (orthogonality)^14,48^. This is particularly important since angles tend to trivially scale with the dimensionality of the data and these methods estimate empirical null distributions for the specific dimensionality being tested. We shuffled one of the vectors under comparison 1000 times by random permutation, and deemed the true angle to be non-orthogonal if it fell left of the 5th percentile of the shuffled distribution.

### Trial-count matching for error decoder analysis

For decoders trained separately on correct or error trials (**Fig. 6a–c, Extended Data Fig. 10b**) we matched the number of trials used to train each decoder using the synthetic minority over-sampling technique (SMOTE^100^). We generated within-distribution synthetic frames by pixel-wise linear interpolation, which preserves the autocorrelation structure of the data. Briefly, for a randomly picked imaging frame *i* at a given spatial position in the maze, we find the *k* nearest-neighbor frames at the same position from other trials (with *k* = 5), pick one of the k frames randomly, *j*, and for each pixel linearly interpolate between *i* and *j*. With this approach, we count-matched correct (*n* = 2539, 40–248 per session) and error (*n* = 1192, 21–109 per session) trials to exactly 300 trials per session, then split into training (80%, *n* = 240 per session) and test (20%, *n* = 60 per session) sets.

### Estimation of spontaneous timescales and related simulations

#### Spontaneous-activity timescales

To estimate spontaneous timescales from across the cortex (**Fig. 4f–i, Extended Data Fig. 9**), we measured widefield Ca^2+^ activity while the mice spontaneously ran in the dark for 20 min in separate sessions that took place before task training. We first removed the potential contributions of running to the activity of each pixel by fitting a linear regression model as described below, separately for each image pixel. We then used the residuals of this model to compute the autocorrelation function of each pixel with a maximum lag of 30 s. The values at non-negative lags of the autocorrelation function *G*(*δ*) were then fitted with both a single and a double exponential, 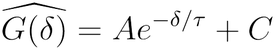 and 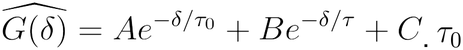. was bounded between 0 and the Nyquist frequency of the signal (i.e., 0.2 s), and *τ* was bounded between the Nyquist frequency and 100 s. In the double exponential fit, *τ*_0_ therefore captures the noise-corrupted part of the autocorrelation, and is considered as the true timescale. We then selected the fit with the highest *R*^*2*^ and defined the spontaneous timescale as the term from the best fitting exponential (see below and **Extended Data Fig. 8** for rationale and further details). We only included pixels with a best-fitting *R*^*2*^ > 0.8 and signal-to-noise ratio (SNR) > 3 (as defined below).

#### Spontaneous-running linear regression

We fit an L2-regularized linear regression model to the spontaneous ΔF/F traces using running speeds at multiple lags as predictors:

Where β’ ‘s are the model coefficients, *τ* ‘s are the lags, *sp*_*t*_ is running speed at time point,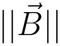 is the L2-norm of the coefficient vector and *λ* is the regularization parameter, determined by cross-validation.

#### SNR

Although the double-exponential method is robust to low SNR in our simulations (**Extended Data Fig. 8**), we excluded low-SNR pixels to further avoid mis-estimation of timescales. We devised a frequency-based SNR metric under the intuition that most signal in widefield data is in low frequencies, whereas most noise lies in higher frequencies after hemodynamic correction. We thus defined 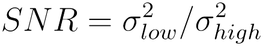, where *low* is ΔF/F low-pass filtered at 3 Hz, *high* is ΔF/F high-pass filtered at 3 Hz (20th-order finite impulse response filters in both cases), and σ^2^ is the variance of the filtered traces.

#### Simulations to validate the exponential-fit-comparison approach

We performed simulations to estimate the robustness of our exponential-fit comparison approach to varying SNR levels. We generated a 20-min time series at 10 Hz (matching our experimental parameters) by drawing from a normal distribution, *x(t) ∼ N*(0,1) . To generate a signal with a single ground-truth timescale *τ*_*true*_, we then convolved *x* with a variably noisy exponential decay filter, 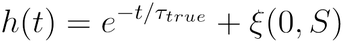, where *S* is systematically scaled to control SNR. We then computed the autocorrelation of the resulting trace and estimated *τ* using either the single or double exponential fits above. The results in **Extended Data Fig. 8c** were obtained using 500 signal simulations for each *τ*_*true*_ at each SNR level. For each value of *S*, the mean effective SNR was calculated from across all simulations as defined above. Importantly, we found that the duration of the trace was crucial to accurately recover *τ*_*true*_ . Specifically, we found that at least 10 minutes of simulated data were necessary to avoid underestimate timescales for *τ*_*true*_ > 10 s (not shown).

#### Simulations of Ca^2+^ dynamics and validation of computation of autocorrelations on behavioral-model residuals

We performed simulations to estimate the impact of (slow) Ca^2+^ indicator dynamics and their behavioral modulation on our ability to estimate *τ*_*true*_ . We first built a model of widefield activity by generating multiple Poisson spiking processes (*N* = 100 per simulation) and simulating their Ca^2+^ binding^101^ and GCaMP6s dynamics^102,103^, and averaging the resulting fluorescence into a single trace. Specifically, we sampled single-unit spikes from a generative firing rate *r*(*t*) and applying a renewal factor as follows. Because directly sampling from a Poisson process results in many refractory period violations, we sampled every *k*-th spiking event from the Poisson (resulting in a gamma instead of exponential distribution inter-spike intervals), markedly reducing these violations. The Ca^2+^ and sensor dynamics were simulated from these spikes using a model of the Ca^2+^ concentration according to the equation:

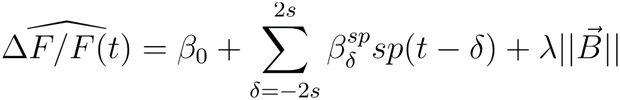

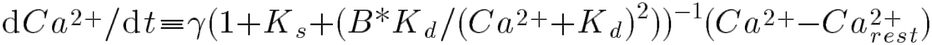

where γ is the Ca^2+^ diffusion constant, is the number of GCaMP6s proteins in the cell being simulated, *K*_*d*_ is the binding affinity, *K*_*s*_ is the binding ratio, and 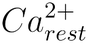 is the baseline free *Ca*^2+^. The modeled Ca^2+^ concentration was then convolved with the double exponential function 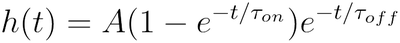 to model the time taken for bound GCaMP6s to become active, where the values of amplitude and time constants *τ*_*on*_ and *τ*_*off*_ were set to capture the specific protein binding dynamics of GCaMP6s. Finally, *ΔF/F* was obtained from the modeled active GCaMP6s sensor by using the Hill equation 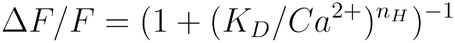, where n_H_ is the Hill coefficient for GCaMP6s. All constants were derived from experimental values found in^102,103^.

| Ca<sub>rest</sub><sup>2+</sup> | γ | K<sub>d</sub> | K<sub>s</sub> | B | A | τ<sub>on</sub> | τ<sub>off</sub> | n<sub>H</sub> |
|---|---|---|---|---|---|---|---|---|
| 50e-9 | 1800 | 147e-9 | 110 | 200e-6 | 7.6e-4 | 0.115 | 1.525 | 2.45 |

To simulate a signal with an intrinsic and a behaviorally driven timescale, we defined *r*(*t*)=*R+n*(*t*)−*b*(*t*), where *R* is the baseline firing rate, *n*(*t*) is activity generated by a process with intrinsic timescale *τ*_*true*_, and *b*(*t*) models a behavioral process with a timescale of *τ*_*behaviour*_ . *n*(*t*) and *b*(*t*) were generated by convolving a Gaussian noise time series with a noiseless exponential decay filter as above (i.e., with *S*=0). Once we obtained a final simulated ΔF/F trace from the average *r*(*t*) over simulated units, we used the same linear regression model as for the data, using lagged values of *b*(*t*) (instead of experimentally measured running speeds) as predictors. We then compared the performance of the double-exponential method applied to autocorrelations computed either on raw *ΔF/F* or residuals of this model, by comparing estimated *τ* with generative *τ*_*true*_over 250 simulations, using a linear correlation coefficient.

### General statistics

For single pairwise comparisons, we used either *t* or signed-rank tests. For comparisons between multiple groups, we applied an analysis of variance (ANOVA, one- or two-way, with or without repeated measures) followed by Tukey’s test for post-hoc comparisons. In situations involving multiple comparisons without ANOVA (e.g., pairwise angle analyses), we used false discovery rate (FDR) correction^104^. Briefly,*n P*-values were ranked in increasing order and the *i-th P*-value (*P*_*i*_) was considered significant at *α* =0.05 if *P*_*i*_*≤(αi)/n*.

## Data availability

The full dataset will be publicly released upon publication of this work.

## Code availability

The source code will be publicly released upon publication of this work.

## Acknowledgements

This work was supported by the US National Institutes of Health (NIH) grants R00MH120047 (to L.P.), R01MH138285 (to L.P.); Simons Foundation grants 872599SPI (to L.P.), NC-GB-PilotExt-00002091 (to L.P.); Alfred P. Sloan Foundation grant SP-2022-19027 (to L.P.); US National Science Foundation grant IOS-2337351 (to L.P.); National Institute for Mathematics and Theory in Biology (Simons Foundations award MP-TMPS-00005320 and National Science Foundation award DMS-2235451) (to L.P.); McKnight Scholar Award (to L.P.); NURTURE postdoctoral fellowship (NIH parent award U54CA272163) (to R.M.C.). We dedicate this manuscript to the memory of our dear friend and colleague Peter Salvino, who passed away during the execution of this work. We thank Julia Cox and David Freedman for thoughtful comments on this manuscript. We also thank Erin Myhre and Lydia VanDeRiet for help training two of the mice, and Xinyue An for her contributions to the development of the SNR metric. Finally, we thank the Brazilian National Council for Scientific and Technological Development for prior support to R.M.C. (CNPq Science without Borders Scholarship 203059/2014-0). Two of the mouse strains used for this research project were obtained from the Mutant Mouse Resource and Research Center (MMRRC), an NIH-funded strain repository. 1) B6(Cg)-Cux2tm2.1(cre)Mull/Mmmh, RRID:MMRRC_032778-MU was donated to the MMRRC at the University of Missouri by Ulrich Mueller, Ph.D., The Scripps Research Institute; 2) B6.FVB(Cg)-Tg(Ntsr1-cre)GN220Gsat/Mmucd, RRID:MMRRC_030648-UCD, was donated by the MMRRC at University of California, Davis, made from the original strain (MMRRC:017266) donated by Nathaniel Heintz, Ph.D., The Rockefeller University, GENSAT and Charles Gerfen, Ph.D., National Institutes of Health, National Institute of Mental Health. Ex-vivo microscopy was performed at the Northwestern University Center for Advanced Microscopy (RRID: SCR_020996) supported by NCI CCSG P30 CA060553 awarded to the Robert H Lurie Comprehensive Cancer Center.

## Author contributions

L.P., R.M.C. and P.S.S. designed the research. L.P., R.M.C., P.S.S. and J.K.L. developed the behavioral task. R.M.C. and J.K.L. collected the imaging data. S.M.I., P.S.S. and R.M.C. trained the mice. S.M.I. performed the histology. L.A.A.-S. developed the timescale-estimation algorithm and performed the simulations. R.M.C. analyzed the data. L.P. and R.M.C. secured the funding. L.P. and R.M.C. wrote the paper and created the data visualizations. L.P. supervised the work.

## Correspondence and requests for materials

should be addressed to Lucas Pinto.

## Extended Data Figures

**Extended Data Fig. 1.**
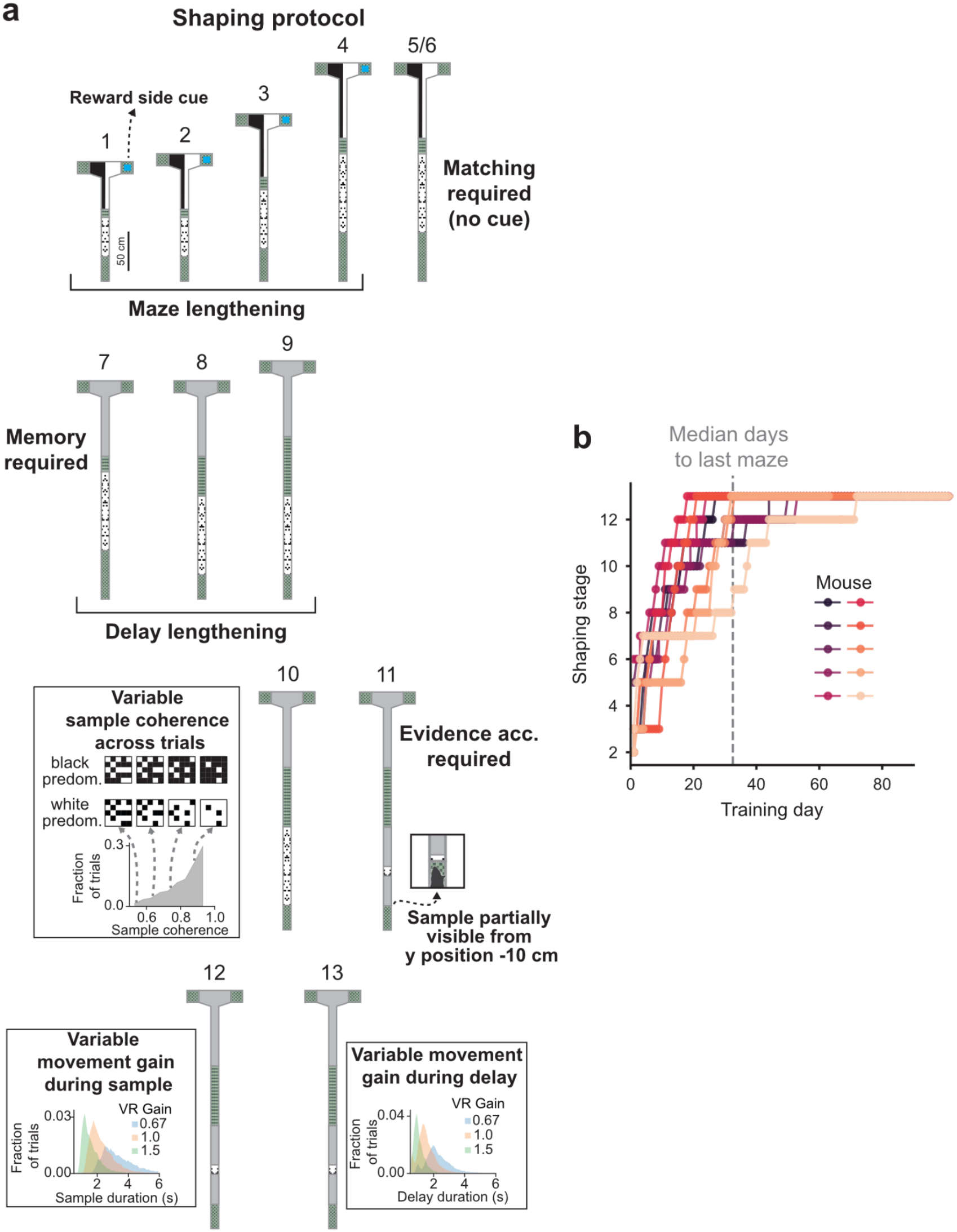
Task shaping. **a**, Maze schematics for all shaping stages. See Methods and **Supplementary Tables 1–4** for more details. **b**, Learning curves for each mouse (colored lines, *N* = 10).

**Extended Data Fig. 2.**
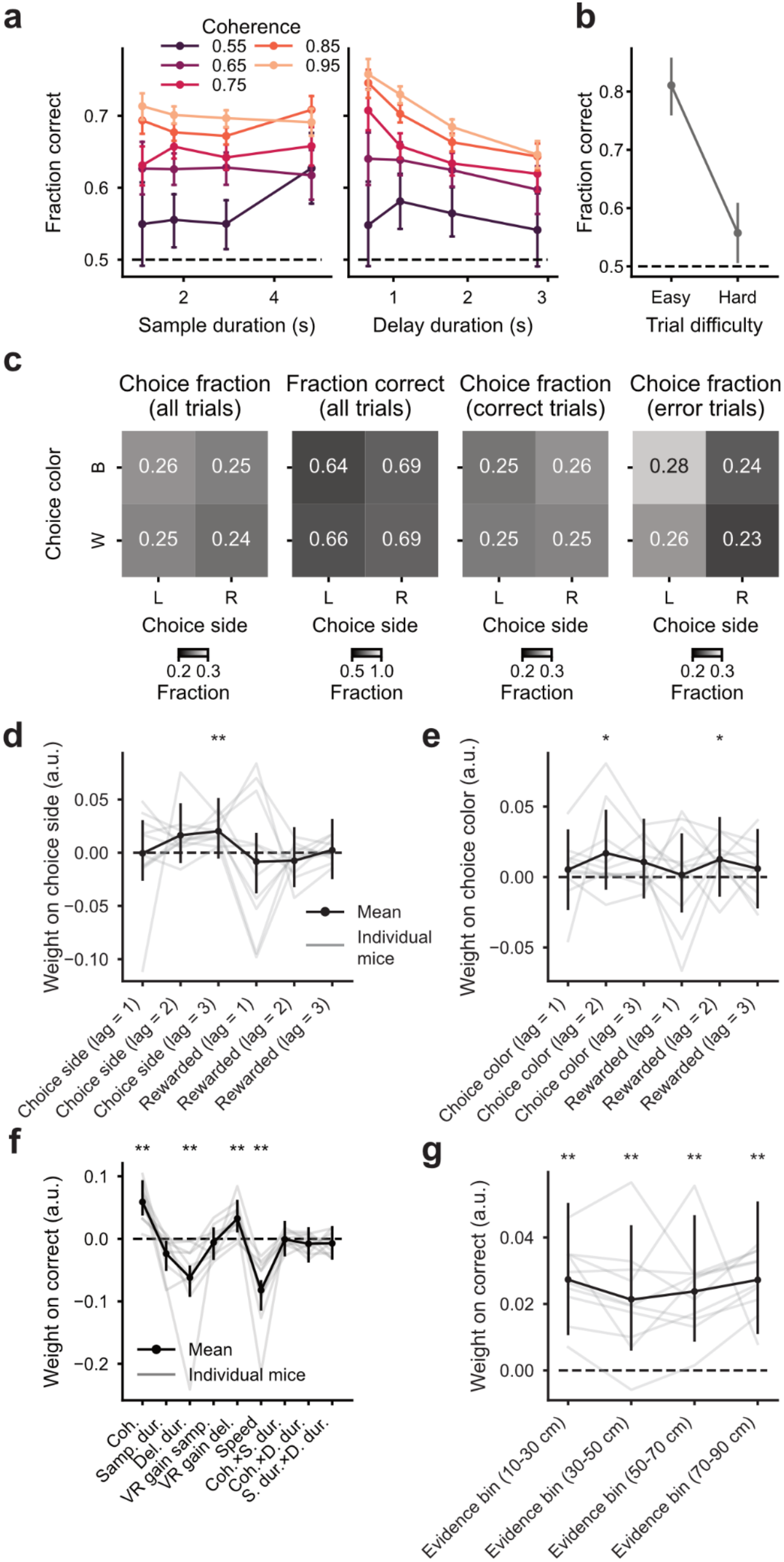
Further quantification of behavioral performance. **a**, Behavioral performance shown separately for different sample coherence levels as a function of sample duration (left) or delay duration (right). Colored lines and datapoints: mean performance for different coherence levels (*n* = 49800 trials across 10 mice). Error bars: 95% confidence intervals (CI), bootstrapped from all trials. **b**, Performance focusing on two regions of the task parameter space where either the easiest or hardest combination of parameters can be found. Easy trials (*n* = 264): high sample coherence (≥ 0.9), long sample duration (3.5–6.0 s), short delay duration (0–1 s). Hard trials (*n* = 384): low sample coherence (0.5–0.65), short sample duration (0–2.5 s), long delay duration (2–4 s). Note that the low number of trials is expected, as the combination of a short sample and a long delay (or vice versa) is least likely when compared to other combinations. Gray datapoints: mean performance. Error bars: 95% CI across trials. **c**, Breakdown of performance and trial fractions for all combinations of chosen side and color, showing minimal bias. L: left, R: right, B: black, W: white. **d**, Coefficients of a logistic regression model trained to predict choice side from trial-history terms. Gray lines: individual mice (*N* = 10). Black data points: average across mice. Error bars: 95% CI bootstrapped per mouse (*n* = 786–7156 trials per mouse) then averaged across mice (*N* = 10). **: *P* < 0.01, *: *P* < 0.05, Wilcoxon signed-rank test against zero with FDR correction. Note that most coefficients are not significantly different from zero, indicating little history bias. **e**, Same as panel d, for chosen color. **f**, Coefficients of a logistic regression model fit to predict the probability of correct choice from the same task parameters as the model in Fig. 1e, with the addition of running speed (*N* = 10 mice). Note that the negative coefficient for running speed indicates that faster running predicts worse performance, whereas the negative coefficient for delay duration indicates that shorter durations predict better performance. Thus, delay duration and running speed affect performance in opposite directions. Conventions as in panel d. **g**, Coefficients from a logistic-regression model trained to predict correct choice from empirical coherence levels from different parts of the sample checkerboard region. Conventions as in panel d. On average, the mice integrate evidence evenly throughout the maze.

**Extended Data Fig. 3.**
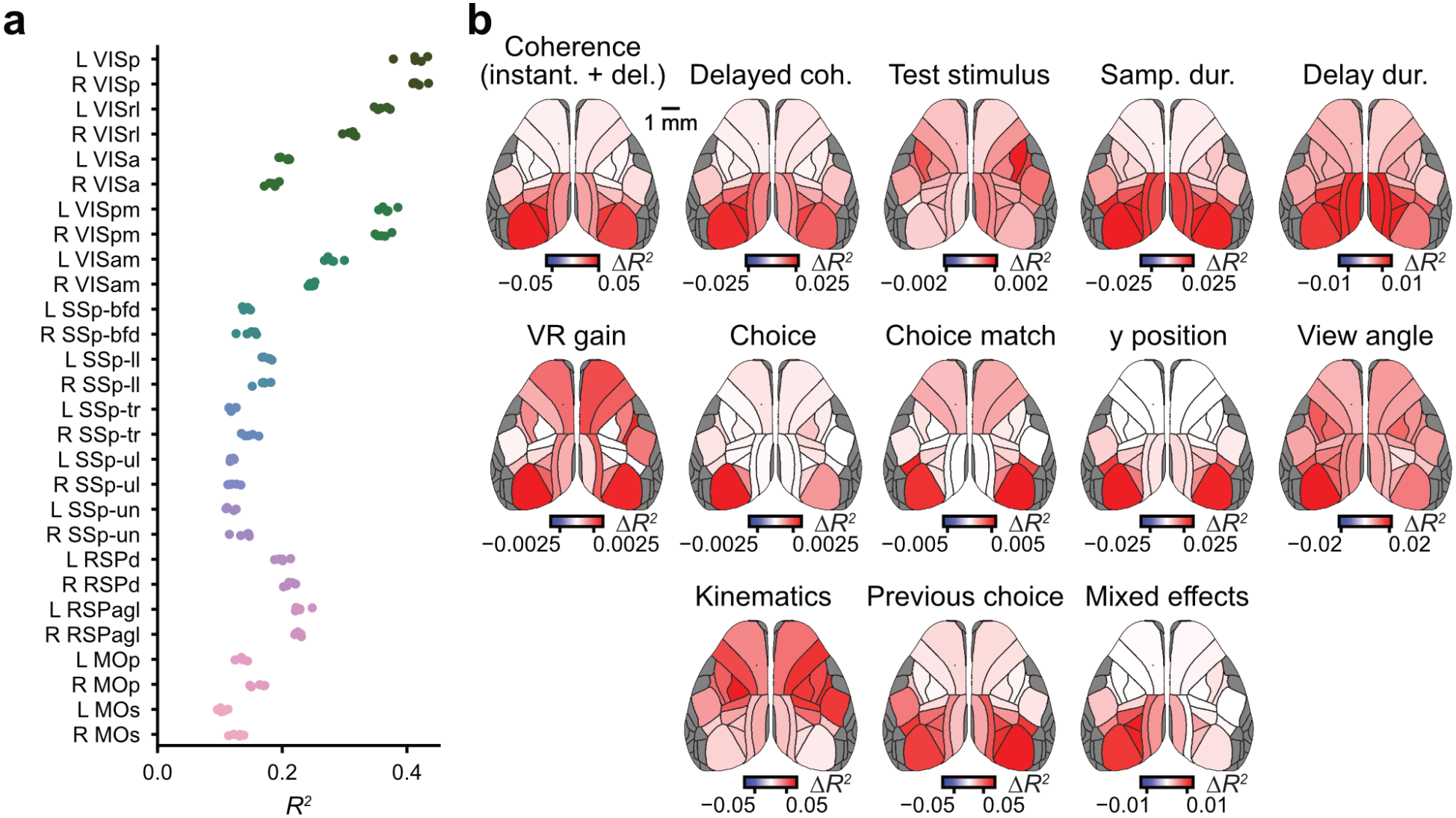
Further quantification of the linear encoding model of neural activity. **a**, Goodness of fit (cross-validated *R*^*2*^) by cortical area. Each model was fitted to data from all sessions (*n* = 4856 trials from 29 sessions and 7 mice). Data points: each of the five data splits. **b**, Cortical-area maps for the change in variance explained when models are refitted without sets of predictors, for all sets. For each map, cross-validated *R*^*2*^ was averaged across five data splits.

**Extended Data Fig. 4.**
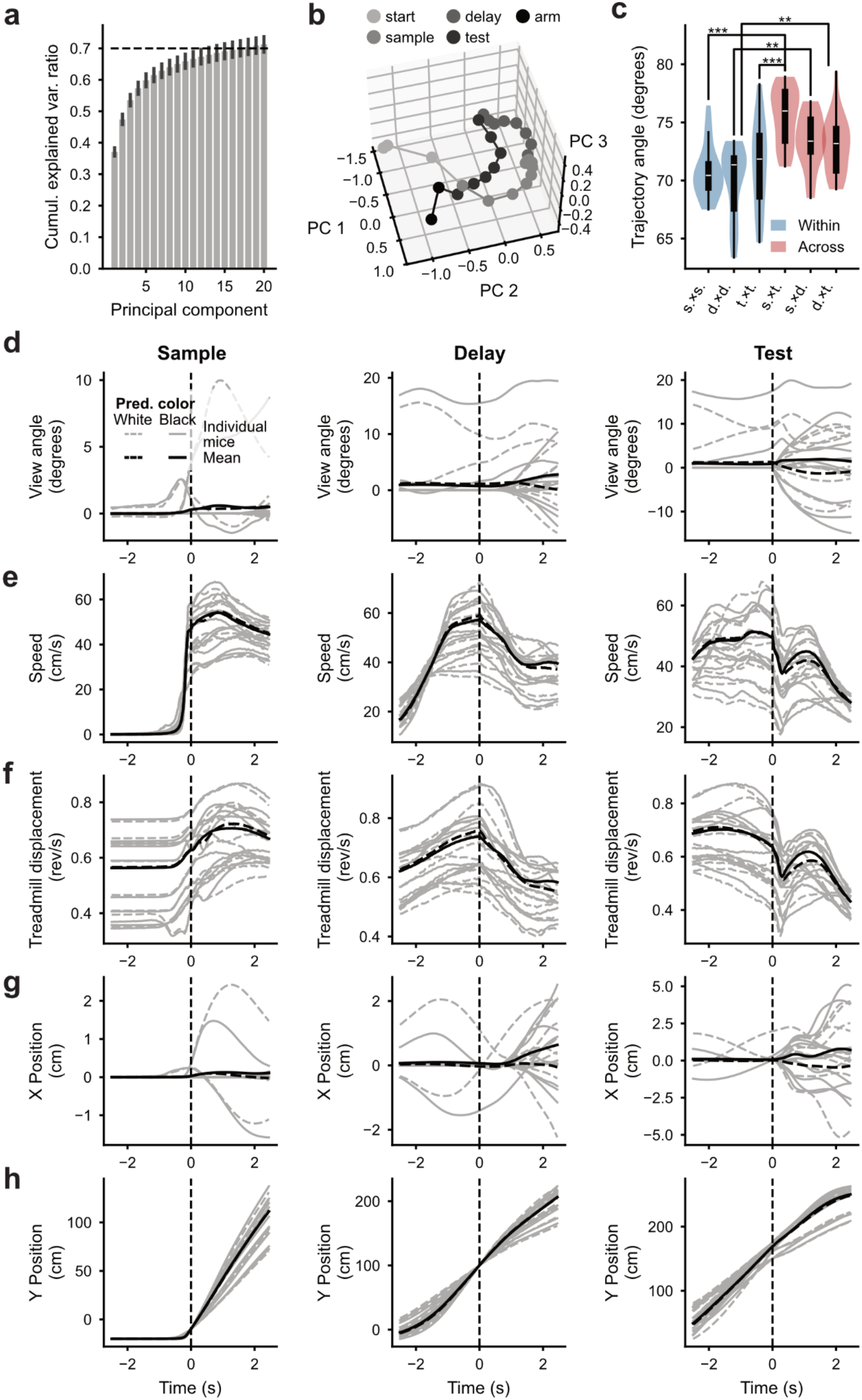
Changes in activity trajectories are not explained by changes in kinematics or luminance alone. **a**, Mean fraction variance explained for the top 20 PCs from the pixel-wise PCA across sessions (*n* = 20 sessions, 5 mice), as in the example session in **Fig. 3a**. Error bars: ± s.e.m.. Dashed line: 70% explained variance. **b**, Average residual activity trajectory projected onto the top three PCs for an example session after regressing out visual scene luminance for the right and left hemifields, position in the virtual maze, running speed, acceleration, treadmill displacement, view angle, and angular velocity. Note that direction changes are preserved in the residual activity trajectory. **c**, Angles between trajectory segments within (blue) vs. across (red) epochs in the residual activity across sessions (*n* = 20 sessions, 5 mice). s.: sample, d.: delay, t.: test. **: *P* < 0.01, ***: *P* < 0.001. Tukey’s post-hoc test after one-way repeated-measures ANOVA. **d**, Virtual view angle over time aligned to the onset of the sample (left), delay (center) or test (right) regions. Gray lines: averages for individual mice across trials (*n* = 786–7156 trials per mouse) shown separately for trials with white predominant sample stimuli (dashed lines, ∼50% of trials) or black predominant stimuli (solid lines). Black lines: averages across mice (*N* = 10). **e–h**, same as **d**, but showing running speed (**e**), treadmill displacement (**f**), x position in the virtual maze (**g**), and y position in the virtual maze (**h**).

**Extended Data Fig. 5.**
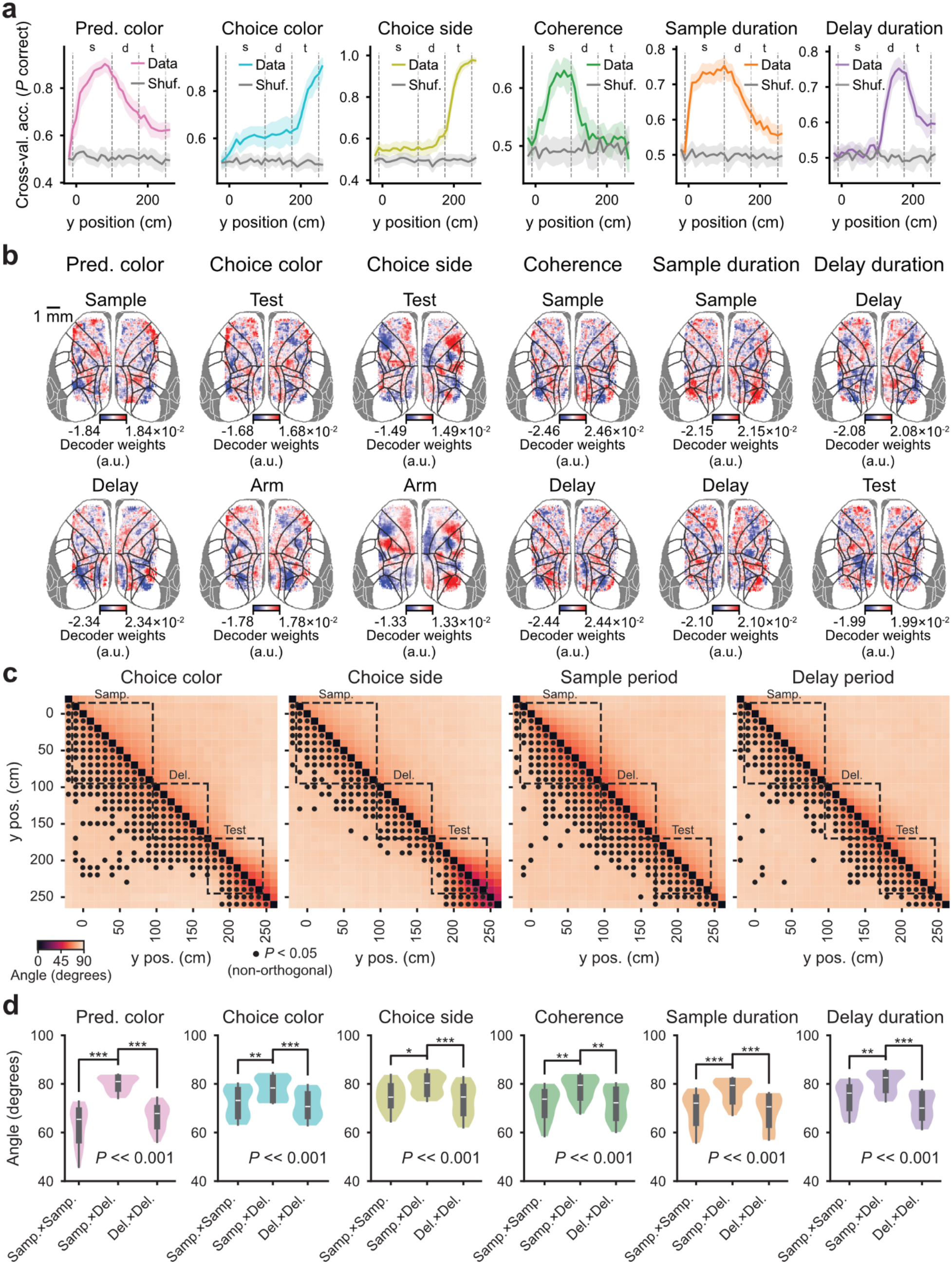
Further quantification of cortex-wide decoders. **a**, Accuracy of binary decoders of task variables from the activity of pixels from across the dorsal cortex, assessed on held-out trials. Different decoders were trained for each session and y position (*n* = 20 sessions, 5 mice). 3731 total trials (67–315 per session) were split into a training set (*n* = 2976, 53–252 trials per set) and a test set (*n* = 755, 14–63 trials per set). Error shades: 95% CI across sessions (*n* = 20). s: sample, d: delay, t: test. **b**, Distributions of decoder weights across the dorsal cortex, for all task variables during relevant epochs, averaged across 20 sessions. **c**, Angles between the decoding axes for the same behavioral variable at different maze positions, for different variables. Dashed boxes: task epochs. Circles: significantly non-orthogonal angles (*P* < 0.05, shuffle test, see Methods for details). Angles and *P*-values are averages across 20 sessions. **d**, Distributions of decoding angles within and across task epochs, for different behavioral variables (*n* = 20 sessions). Printed *P*-value is from a one-way repeated-measures ANOVA with factor ‘epoch pairs’. *: *P* < 0.05, **: *P* < 0.01, ***: *P* < 0.001, Tukey’s post-hoc test. Note that, for all variables, angles are significantly more orthogonal across than within epochs.

**Extended Data Fig. 6.**
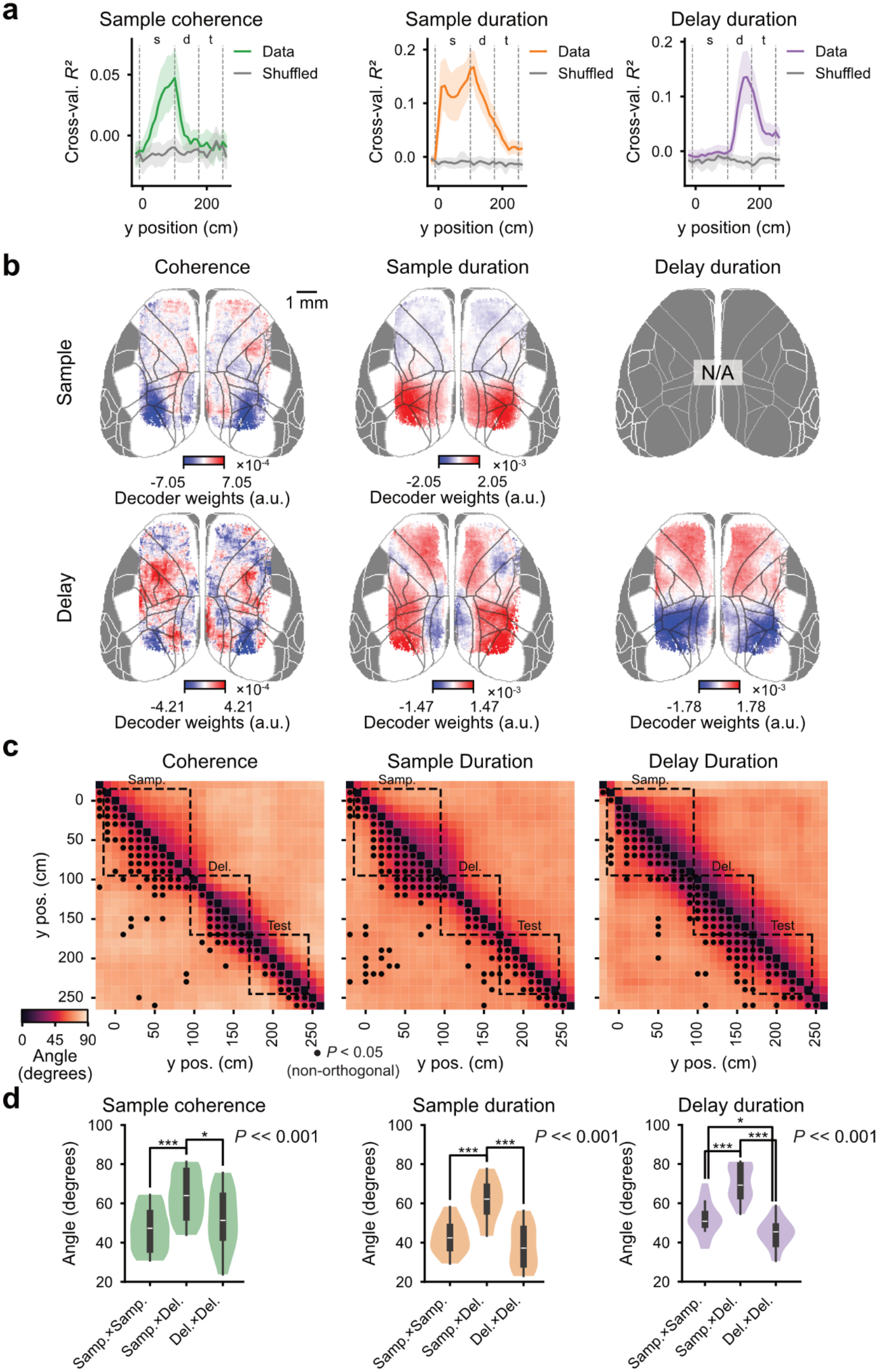
Regression-based decoders for continuous variables also capture dynamic coding subspaces. **a**, Cross-validated performance of cortex-wide regression-based SVM decoders of continuous task variables (coherence, sample duration, and delay duration). Different decoders were trained for each y position. *R*^*2*^ is from 755 held-out trials. Note that cross-validated *R*^*2*^ can dip below zero because the training and test distributions are not identical. Error shades: 95% CI across sessions (*n* = 20). s: sample, d: delay, t: test. **b**, Distributions of decoder weights across the dorsal cortex, for all task variables during sample and delay epochs, averaged across 20 sessions. **c**, Angles between the decoding axes for the same behavioral variable at different maze positions, for different variables. Dashed boxes: task epochs. Circles: significantly non-orthogonal angles (*P* < 0.05, shuffle test, see Methods for details). Angles and *P*-values are averages across 20 sessions. Note that, due to being prone to overfitting, these models required strong L2 regularization, which brings weights closer to zero, therefore likely overestimating the degree of collinearity between models. **d**, Distributions of decoding angles within and across task epochs, for different behavioral variables (*n* = 20 sessions). Printed *P*-value is for a one-way repeated-measures ANOVA with factor ‘epoch pairs’. *: *P* < 0.05, **: *P* < 0.01, ***: *P* < 0.001, Tukey’s post-hoc test. Note that, for all variables, angles are significantly more orthogonal across than within epochs, as is the case with binarized classifiers (**Fig. 3, Extended Data Fig. 5**).

**Extended Data Fig. 7.**
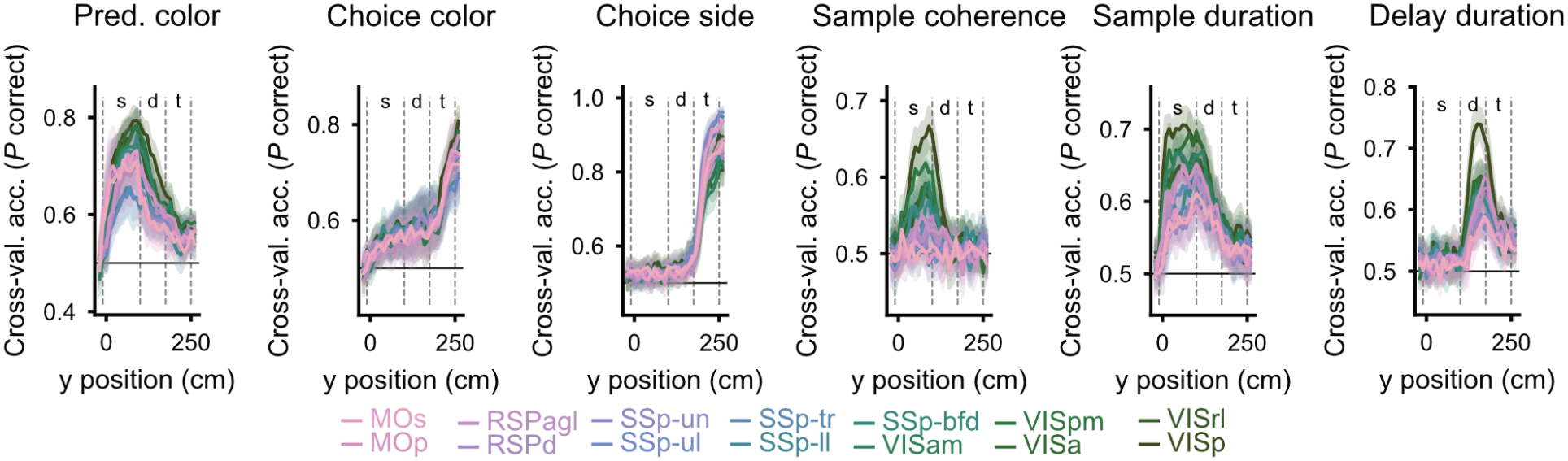
Area-specific decoders confirm distributed task information throughout the cortex. Accuracy of decoders for each behavioral variable, trained on pixels taken separately from each cortical region, assessed on held-out trials. Different decoders were trained for each session, cortical region and y position (*n* = 20 sessions, 5 mice). 3731 total trials (67–315 per session) were split into a training set (*n* = 2976, trials per 53–252 per set) and a testing set (*n* = 755, 14–63 trials per set). Error shades: 95% CI across 20 sessions. s: sample, d: delay, t: test. The ‘predominant color’ decoders are the same as in **Fig. 4a**, reproduced here for convenience.

**Extended Data Fig. 8.**
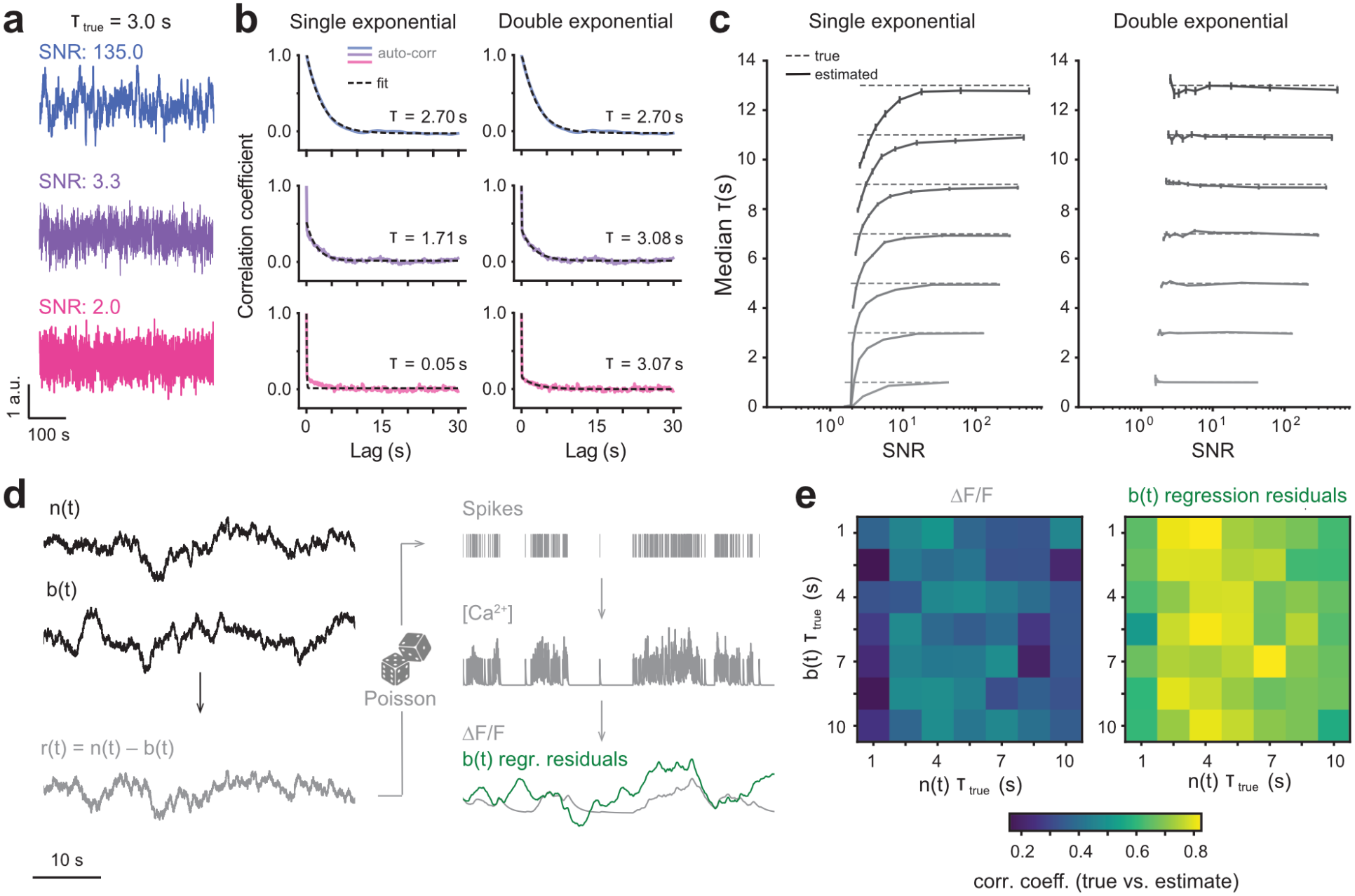
Simulations to validate our method for spontaneous-timescale estimation (see Methods for details). **a**, Example traces generated with the same timescale (by convolving a random Gaussian vector with an exponential decay kernel), and corrupted with increasing levels of Gaussian noise. **b**, Example single and double exponential fits for the traces in panel a. **c**, Median ± s.e.m. of estimated timescale across 500 simulations as in panels a, b, for different generative timescales (indicated by the dashed lines), and for single vs. double exponential fits. **d**, Schematic of the simulation of Ca^2+^ signals from spiking data with underlying rates of known timescale, *n(t*), linearly combined with another signal *b(t*). In these simulations, the goal is to regress out *b(t*) to estimate the underlying timescale of *n(t*). **e**, Correlation between true and estimated timescale of *n(t*), computed over 250 simulations, computed using either the simulated ΔF/F or the residuals from the *b(t*) regression.

**Extended Data Fig. 9.**
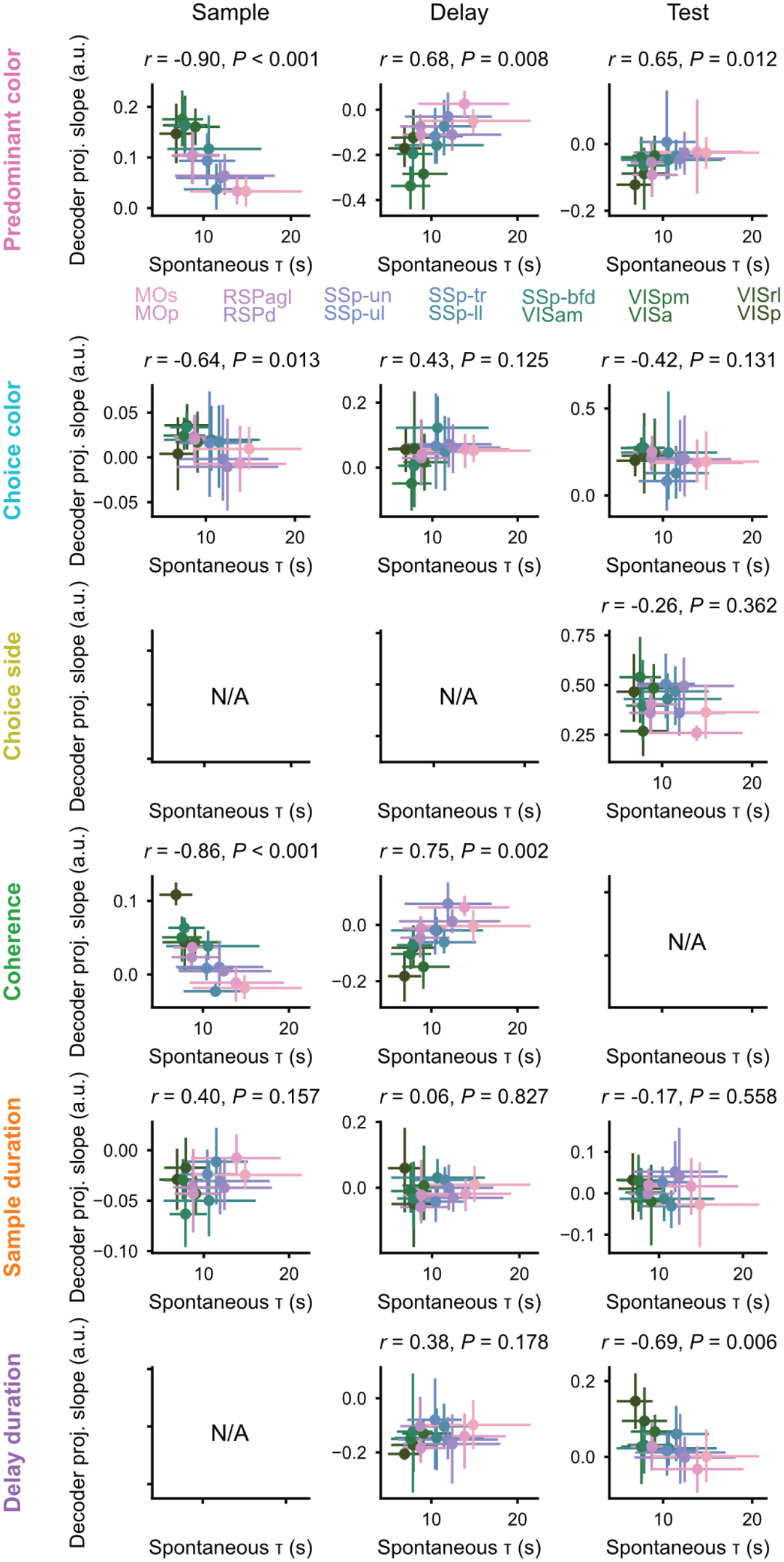
Correlations between spontaneous activity timescales and activity projections for other behavioral variables. Scatter plots of average area timescale vs. the average slope of projected activity in the coding subspaces for each behavioral variable during the sample, delay, and test epochs. For each area, slopes from 755 held-out trials (14–63 per session) across 20 sessions (3–6 sessions per mouse) were averaged by mouse (*N* = 5). Error bars: 95% CI across 5 mice.

**Extended Data Fig. 10.**
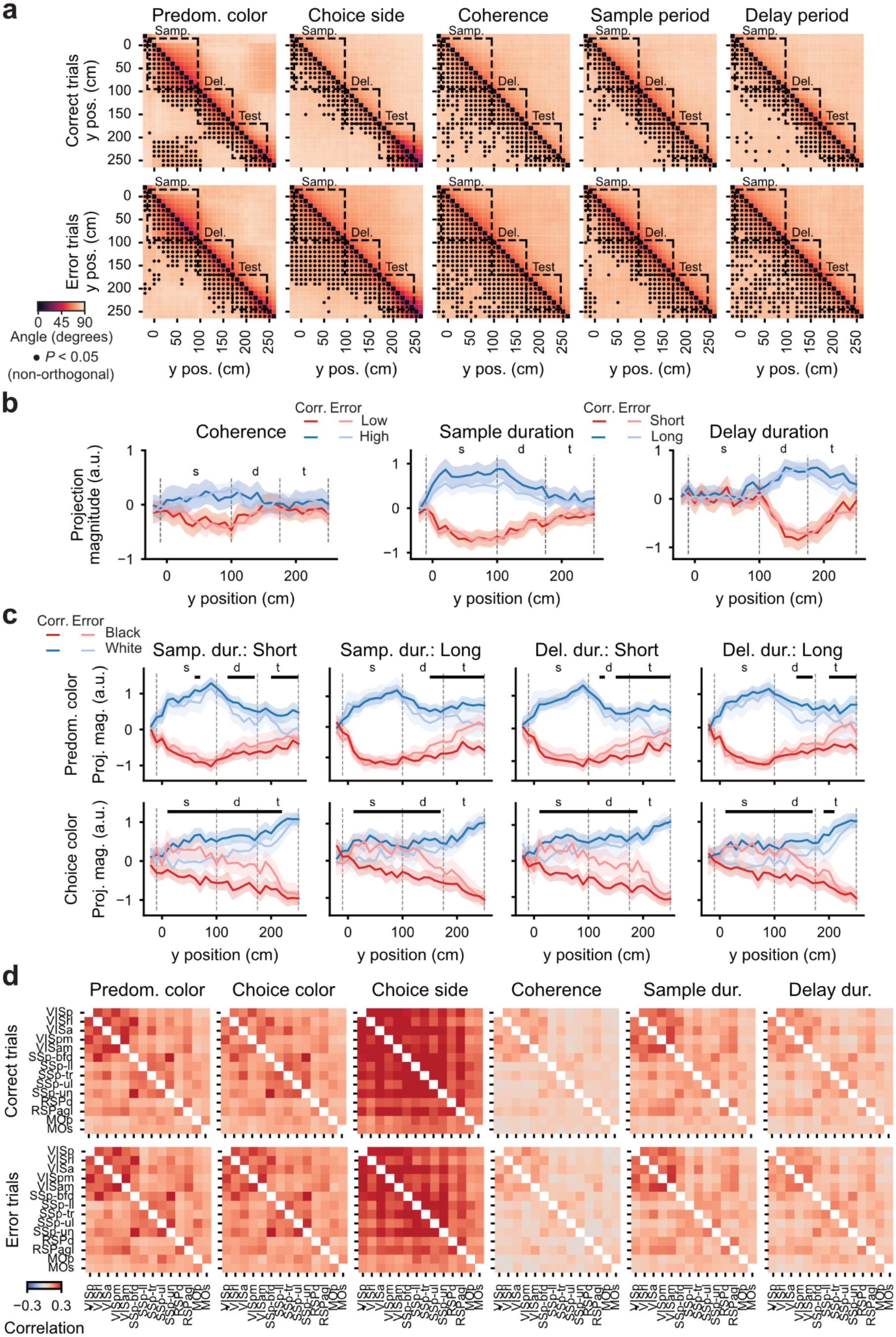
Error trials are associated with less orthogonal angles. **a**, Angles between the decoding axes for the same behavioral variable at different maze positions, for different variables, plotted separately for correct and error trials. Dashed boxes: task epochs. Circles: significantly non-orthogonal angles (*P* < 0.05, shuffle test). **b**, Cortex-wide activity projected onto the ‘sample coherence’, ‘sample duration’, and ‘delay duration’ decoding axes, separately for correct and error trials, and combined across sessions (*n* = 494 correct, 261 error held-out trials). Error shades: 95% CI across trials. There were no significant differences in projection magnitude between correct and error trials (*P* > 0.05, permutation test). **c**, Cortex-wide activity during trials with short or long sample duration (left two plots) or delay duration (right two plots), projected onto the ‘predominant color’ and ‘choice color’ decoding axes separately for correct and error trials (*n* = 494 correct, 261 error held-out trials, split along the median for either sample or delay duration). Error shades: 95% CI across trials. Horizontal bars on top: *P* < 0.05, permutation test (correct vs. error trials, combined for black and white predominant or chosen color). **d**, Pairwise correlation between single-area decision variables for different task parameters, shown separately for correct and error trials (*n* = 494 correct, 261 error held-out trials).

## Supplementary Figures

**Supplementary Fig. 1.**
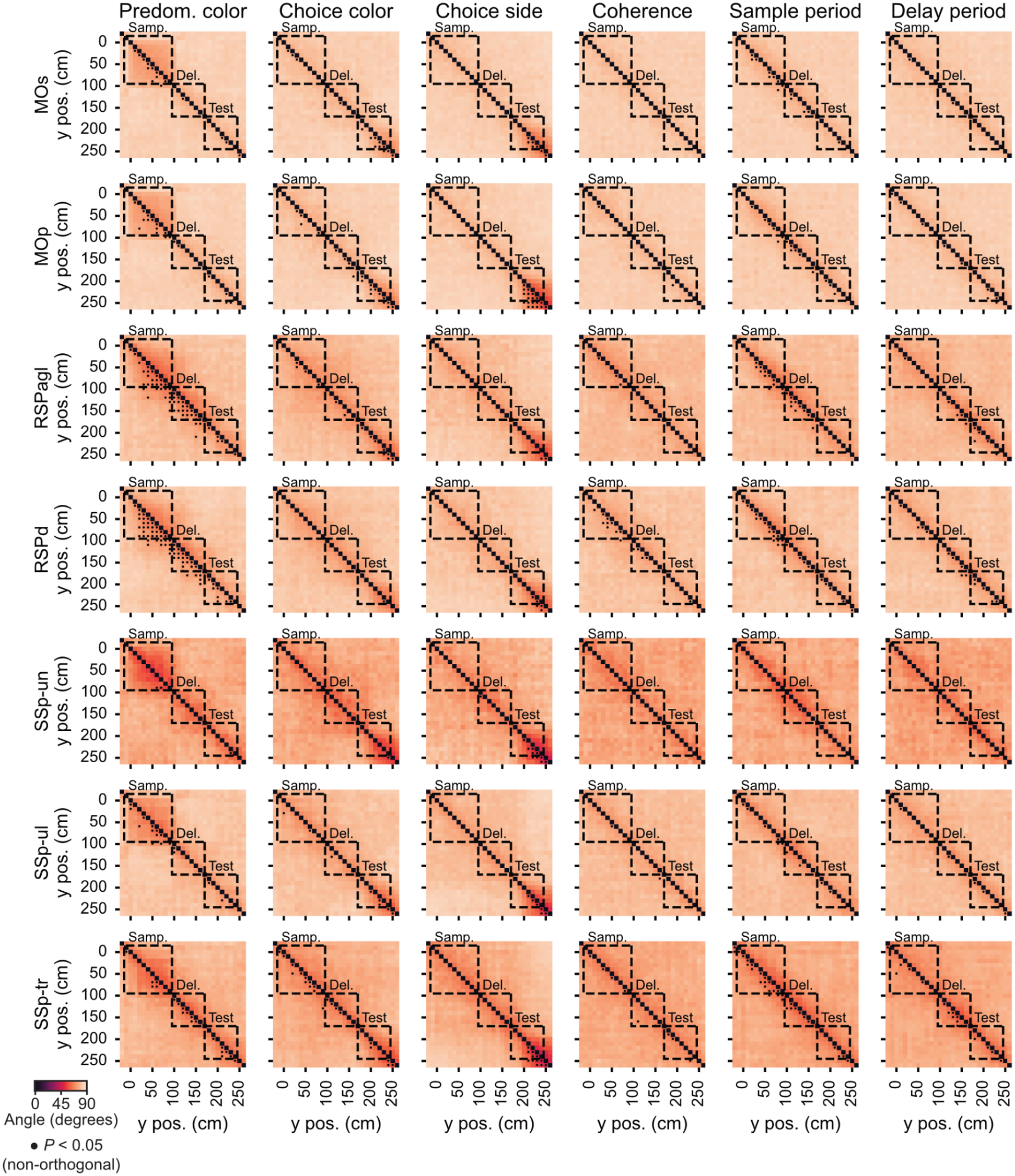
Dynamic cognitive encoding within cortical areas. Decoders were trained on pixels from single anatomically defined cortical areas. Shown are the angles between the decoding axes for the same behavioral variable at different maze positions, for different variables. Dashed boxes: task epochs. Circles: significantly non-orthogonal angles (*P* < 0.05, shuffle test). Angles and *P*-values are averages across 20 sessions.

**Supplementary Fig. 2.**
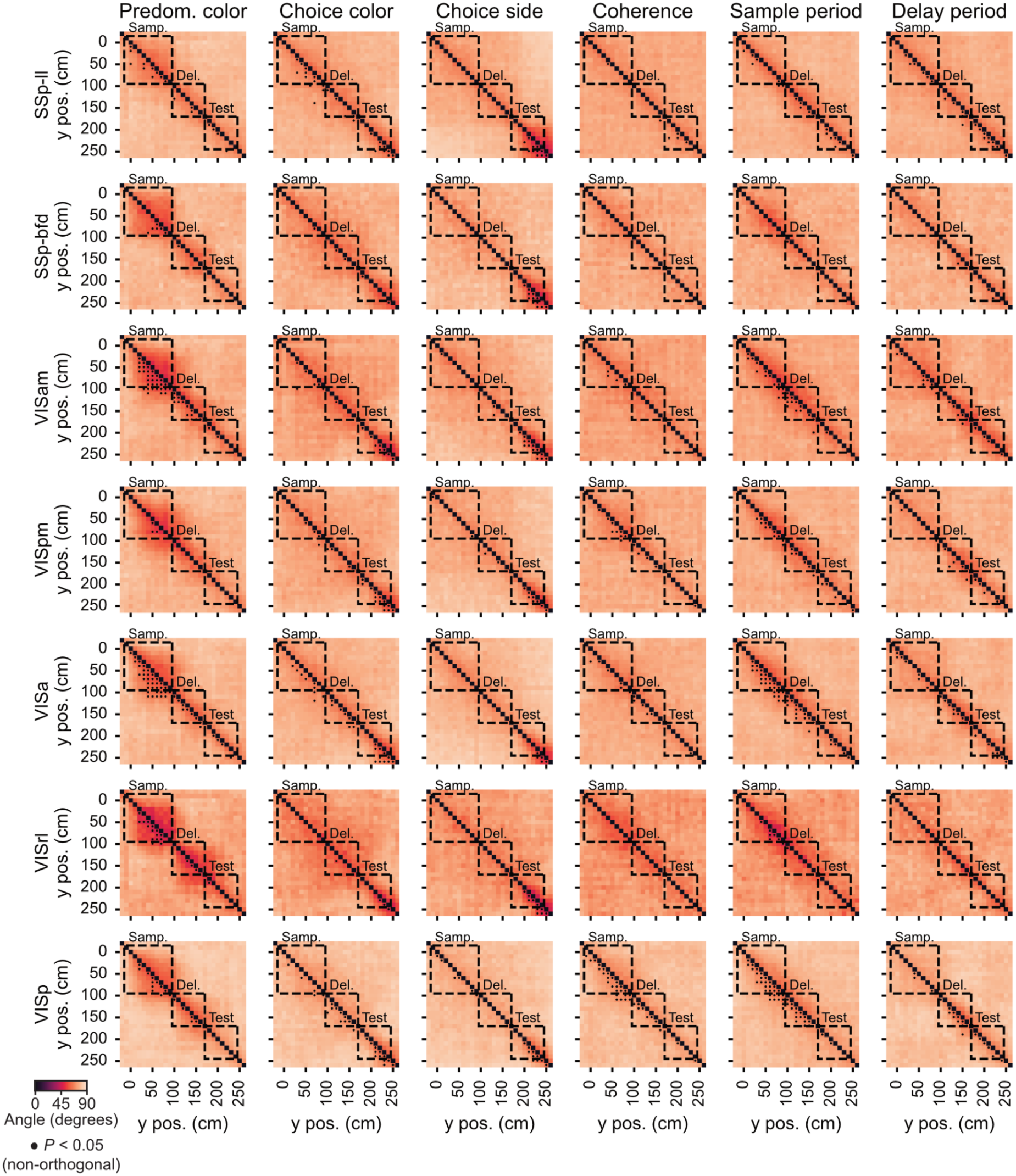
Dynamic cognitive encoding within cortical areas. Same as **Supplementary Fig. 1**, for the other imaged cortical areas.

**Supplementary Fig. 3.**
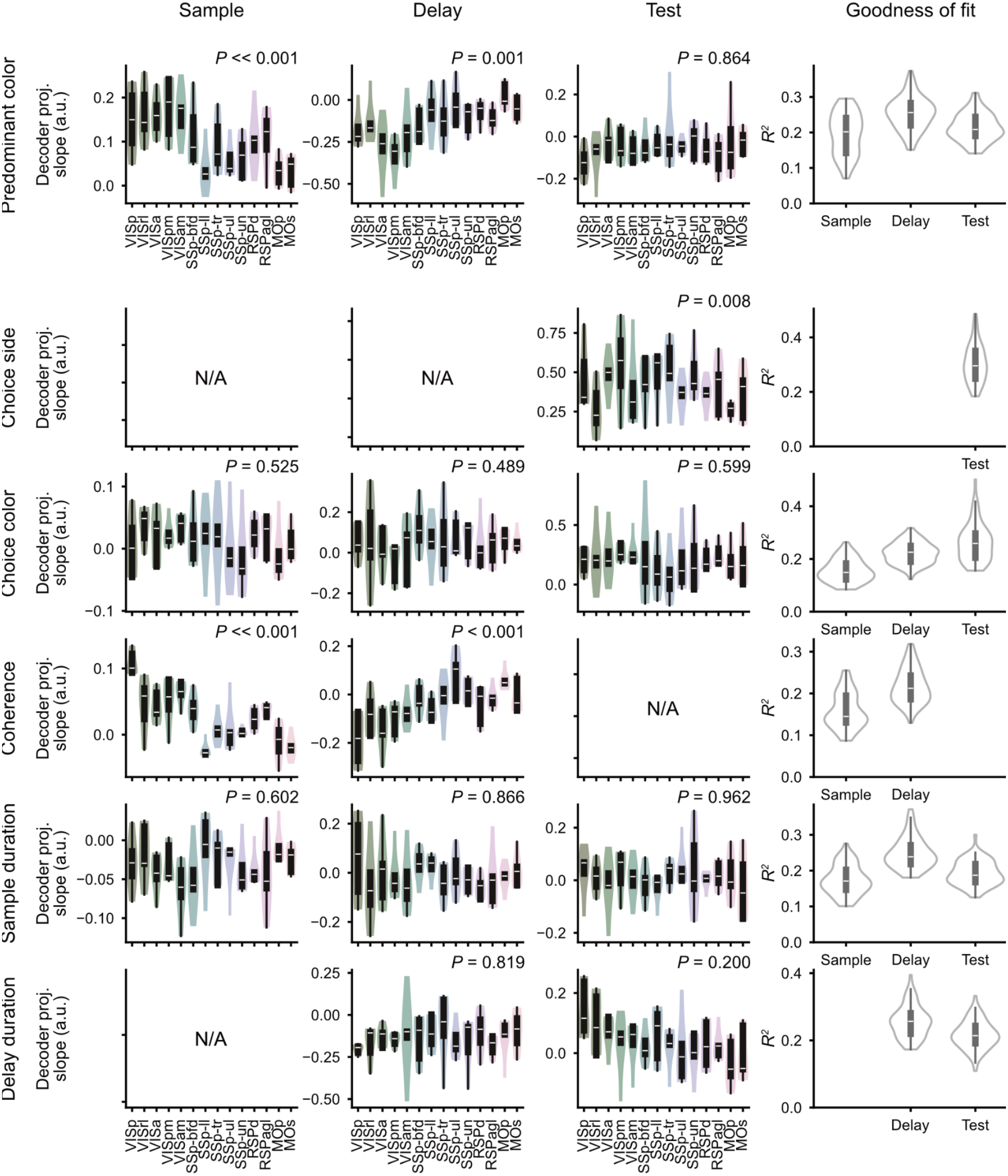
Slopes of linear fits to activity projections across region-specific coding subspaces for different behavioral variables and task epochs. *First three columns from the left*: unsigned slope distributions across cortical regions of linear fits during sample, delay, and test epochs for activity projected onto the decoding axes for each behavioral variable. Printed *P*-value: one-way repeated-measures ANOVA. *Rightmost column*: goodness of linear fit to projected-activity time courses during each task epoch.

## Supplementary Tables

**Supplementary Table 1.** Maze y dimensions for the different shaping steps. Dimensions in cm for different maze regions. The start of the sample checkerboard region is y = 0 by convention.

| **Maze y dimensions** |  |  |  |  |  |  |  |  |  |
|---|---|---|---|---|---|---|---|---|---|
| **Shaping step** | **Maze ID** | **Step description** | **Start position** | **Sample start** | **Delay start** | **Test start** | **Choice arm start** | **Reward zone start** | **Maze end** |
| **1** | M1 | Cued reward | -15 | 0 | 50 | 60 | 100 | 105 | 120 |
| **2** | M2 | Maze lengthening<br>(cued reward) | -15 | 0 | 50 | 60 | 110 | 115 | 130 |
| **3** | M3 | Maze lengthening<br>(cued reward) | -30 | 0 | 75 | 90 | 150 | 155 | 170 |
| **4** | M4 | Maze lengthening<br>(cued reward) | -50 | 0 | 100 | 120 | 195 | 200 | 215 |
| **5** | M8 | Match rule<br>(interspersed cued trials) | -50 | 0 | 100 | 120 | 195 | 200 | 215 |
| **6** | M10 | Match rule<br>(no cued trials) | -50 | 0 | 100 | 120 | 195 | 200 | 215 |
| **7** | M11 | Memory<br>requirement | -50 | 0 | 100 | 120 | 195 | 200 | 215 |
| **8** | M13 | Memory<br>requirement<br>(delay lengthening) | -20 | 0 | 100 | 150 | 225 | 230 | 245 |
| **9** | M15 | Memory<br>requirement<br>(delay lengthening) | -20 | 0 | 100 | 175 | 250 | 255 | 270 |
| **10** | M17 | Variable sample<br>coherence | -20 | 0 | 100 | 175 | 250 | 255 | 270 |
| **11** | M18 | Evidence<br>accumulation<br>requirement | -20 | 0 | 100 | 175 | 250 | 255 | 270 |
| **12** | M19 | Variable sample<br>VR gain | -20 | 0 | 100 | 175 | 250 | 255 | 270 |
| **13** | M20 | Variable delay<br>VR gain | -20 | 0 | 100 | 175 | 250 | 255 | 270 |

**Supplementary Table 2.** Important task parameters for the different shaping steps. Reward cue is a visual guide indicating the rewarded arm. A masked test stimulus is one that is revealed only when the mouse reaches the test region. Sample visibility refers to the width of the checkerboard sliver the mouse sees as it runs. An infinite value indicates that the full checkerboard is visible.

| **Task parameters** |  |  |  |  |  |  |  |  |
|---|---|---|---|---|---|---|---|---|
| **Shaping step** | **Reward cue** | **Masked test** | **Sample visibility** | **Variable sample coherence** | **Variable sample gain** | **Variable delay gain** | **Sample coherence distribution** | **VR gains** |
| **1** | Yes | No | Infinite | No | No | No | Constant 0.95 coherence | 1 |
| **2** | Yes | No | Infinite | No | No | No | Constant 0.95 coherence | 1 |
| **3** | Yes | No | Infinite | No | No | No | Constant 0.95 coherence | 1 |
| **4** | Yes | No | Infinite | No | No | No | Constant 0.95 coherence | 1 |
| **5** | No | No | Infinite | No | No | No | Constant 0.95 coherence | 1 |
| **6** | No | No | Infinite | No | No | No | Constant 0.95 coherence | 1 |
| **7** | No | Yes | Infinite | No | No | No | Constant 0.95 coherence | 1 |
| **8** | No | Yes | Infinite | No | No | No | Constant 0.95 coherence | 1 |
| **9** | No | Yes | Infinite | No | No | No | Constant 0.95 coherence | 1 |
| **10** | No | Yes | Infinite | Yes | No | No | Exponential | 1 |
| **11** | No | Yes | 20 cm | Yes | No | No | Exponential | 1 |
| **12** | No | Yes | 20 cm | Yes | Yes | No | Exponential | 0.67 (30%), 1 (40%), 1.5 (30%) |
| **13** | No | Yes | 20 cm | Yes | Yes | Yes | Exponential | 0.67 (30%), 1 (40%), 1.5 (30%) |

**Supplementary Table 3.**
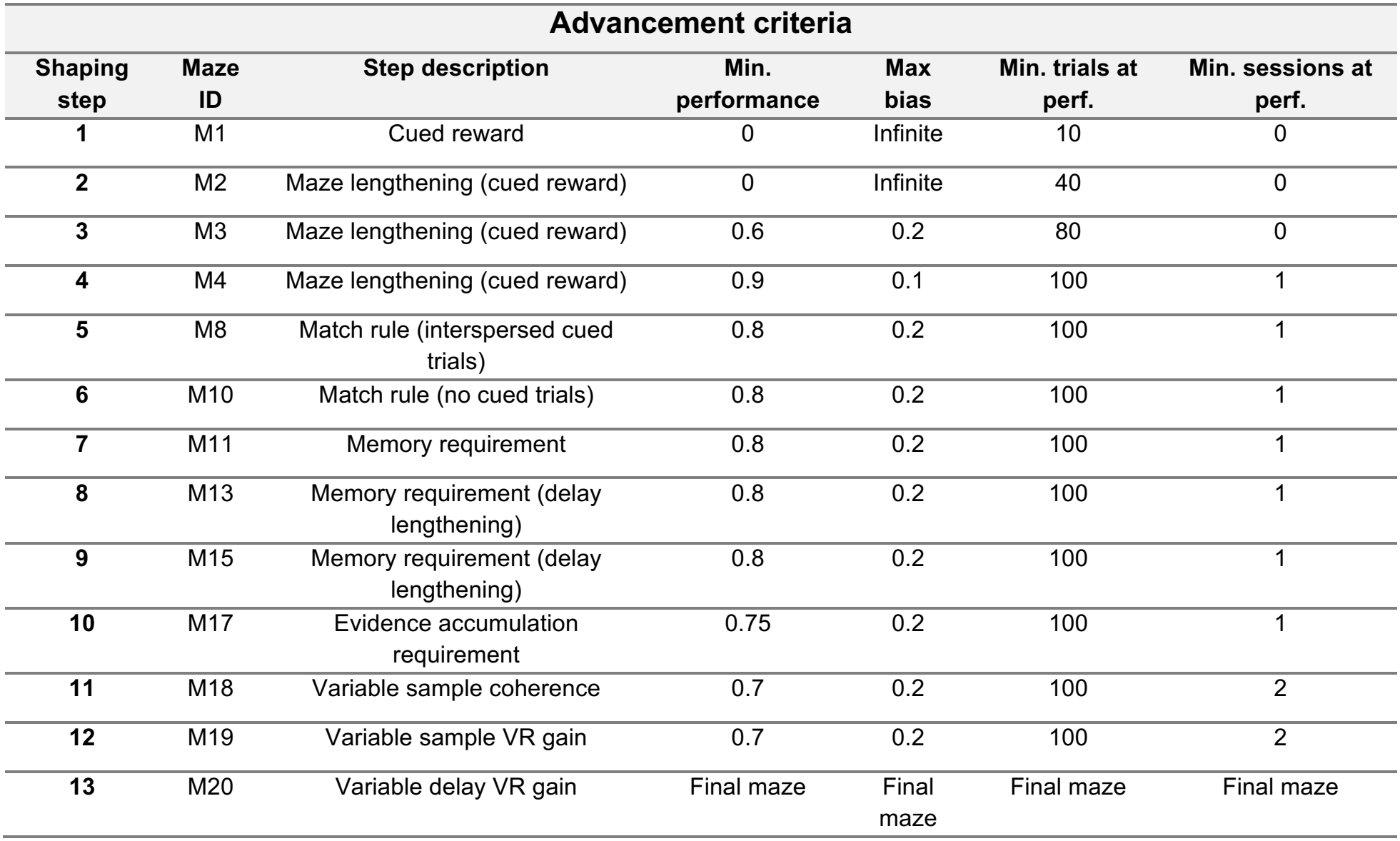
Advancement criteria for the different shaping steps. For each shaping step, we show the number of consecutive trials at criterion (minimum proportion correct and maximum side bias) required to automatically advance to the next stage.

| **Advancement criteria** |  |  |  |  |  |  |
|---|---|---|---|---|---|---|
| **Shaping step** | **Maze ID** | **Step description** | **Min. performance** | **Max bias** | **Min. trials at perf.** | **Min. sessions at perf.** |
| **1** | M1 | Cued reward | 0 | Infinite | 10 | 0 |
| **2** | M2 | Maze lengthening (cued reward) | 0 | Infinite | 40 | 0 |
| **3** | M3 | Maze lengthening (cued reward) | 0.6 | 0.2 | 80 | 0 |
| **4** | M4 | Maze lengthening (cued reward) | 0.9 | 0.1 | 100 | 1 |
| **5** | M8 | Match rule (interspersed cued trials) | 0.8 | 0.2 | 100 | 1 |
| **6** | M10 | Match rule (no cued trials) | 0.8 | 0.2 | 100 | 1 |
| **7** | M11 | Memory requirement | 0.8 | 0.2 | 100 | 1 |
| **8** | M13 | Memory requirement (delay lengthening) | 0.8 | 0.2 | 100 | 1 |
| **9** | M15 | Memory requirement (delay lengthening) | 0.8 | 0.2 | 100 | 1 |
| **10** | M17 | Evidence accumulation requirement | 0.75 | 0.2 | 100 | 1 |
| **11** | M18 | Variable sample coherence | 0.7 | 0.2 | 100 | 2 |
| **12** | M19 | Variable sample VR gain | 0.7 | 0.2 | 100 | 2 |
| **13** | M20 | Variable delay VR gain | Final maze | Final maze | Final maze | Final maze |

**Supplementary Table 4.** Easy-trial block information for the different shaping steps. In most mazes, the mice occasionally transition into a short block of easy trials to help maintain task engagement, with maze-dependent block lengths and block-transition triggers (random vs. performance-based).

| **Easy-trial block information** |  |  |  |  |  |  |
|---|---|---|---|---|---|---|
| **Shaping step** | **Maze ID** | **Step description** | **Easy maze ID** | **Trigger type** | **Number of easy trials** | **Min. performance** |
| **1** | M1 | Cued reward | None | None | 0 | 0 |
| **2** | M2 | Maze lengthening (cued reward) | None | None | 0 | 0 |
| **3** | M3 | Maze lengthening (cued reward) | None | None | 0 | 0 |
| **4** | M4 | Maze lengthening (cued reward) | None | None | 0 | 0 |
| **5** | M8 | Match rule (interspersed cued trials) | M4 | Randomly interspersed | 5–15 | 0 |
| **6** | M10 | Match rule (no cued trials) | None | None | 0 | 0 |
| **7** | M11 | Memory requirement | M10 | Randomly interspersed | 5–15 | 0 |
| **8** | M13 | Memory requirement (delay lengthening) | M11 | Randomly interspersed | 5–15 | 0 |
| **9** | M15 | Memory requirement (delay lengthening) | M13 | Randomly interspersed | 5–15 | 0 |
| **10** | M17 | Evidence accumulation requirement | M4 | Performance triggered | 10 | 0.55 |
| **11** | M18 | Variable sample coherence | M4 | Performance triggered | 10 | 0.55 |
| **12** | M19 | Variable sample VR gain | M4 | Performance triggered | 10 | 0.55 |
| **13** | M20 | Variable delay VR gain | M4 | Performance triggered | 10 | 0.55 |

**Supplementary Table 5.**
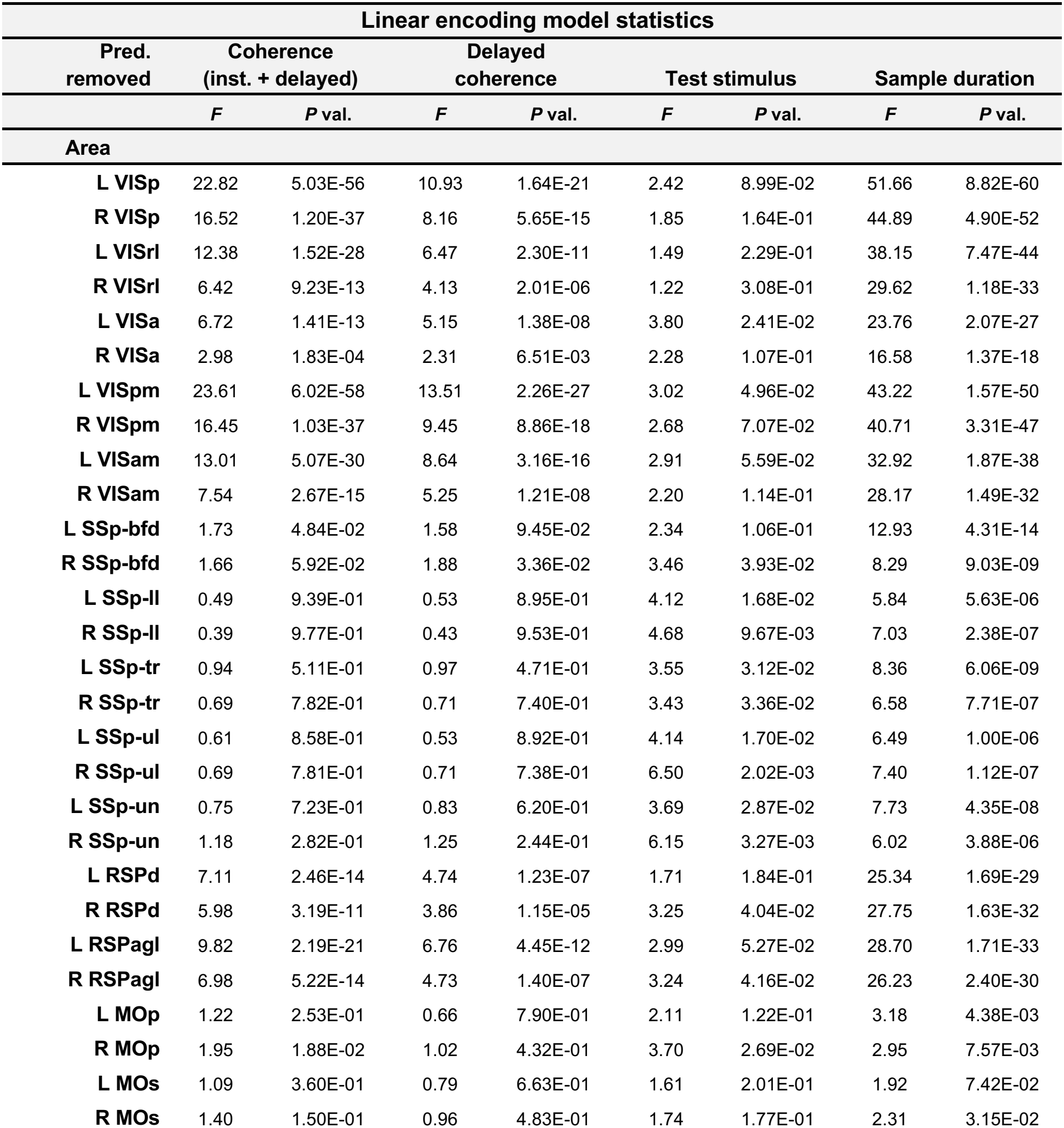
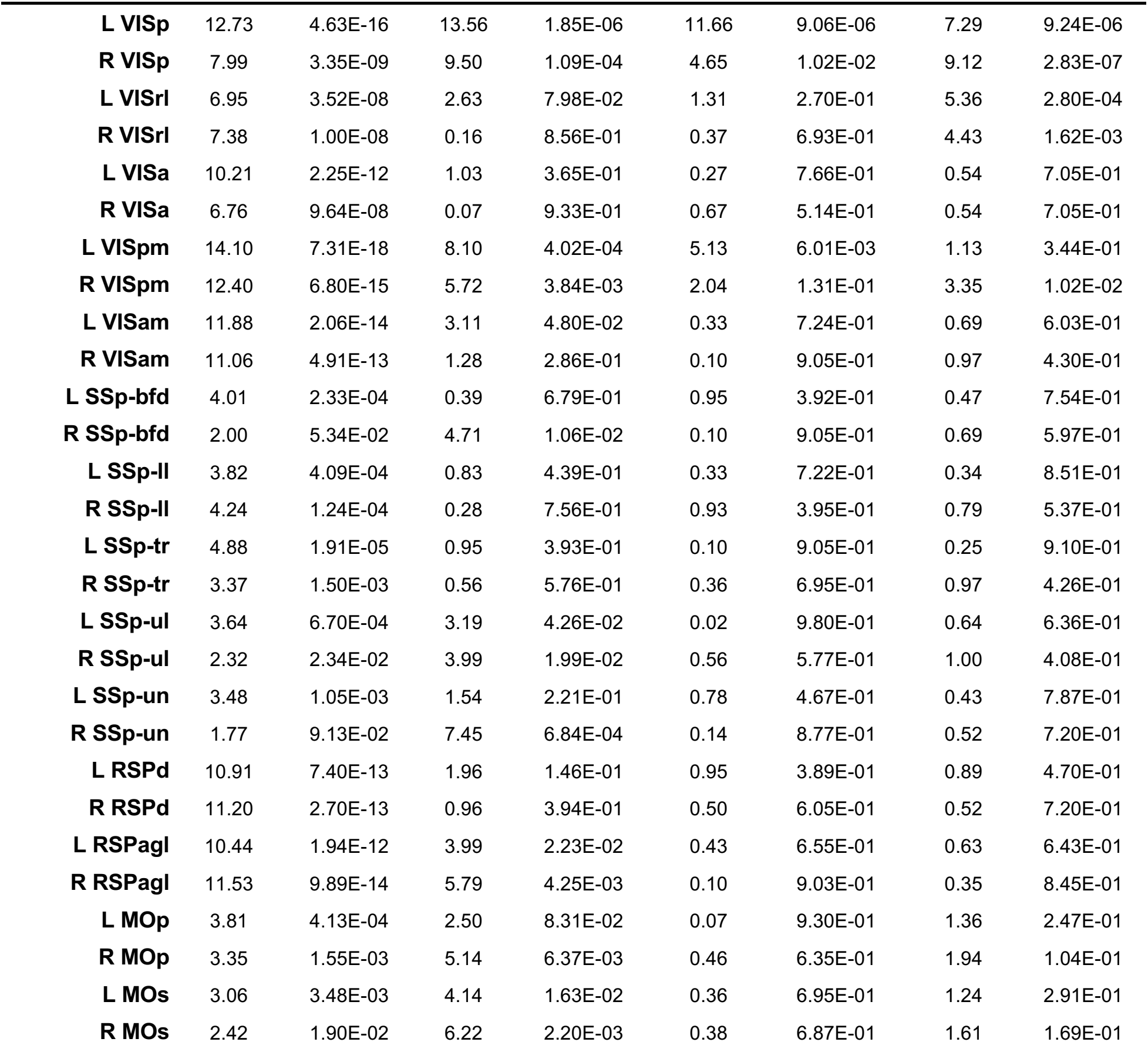
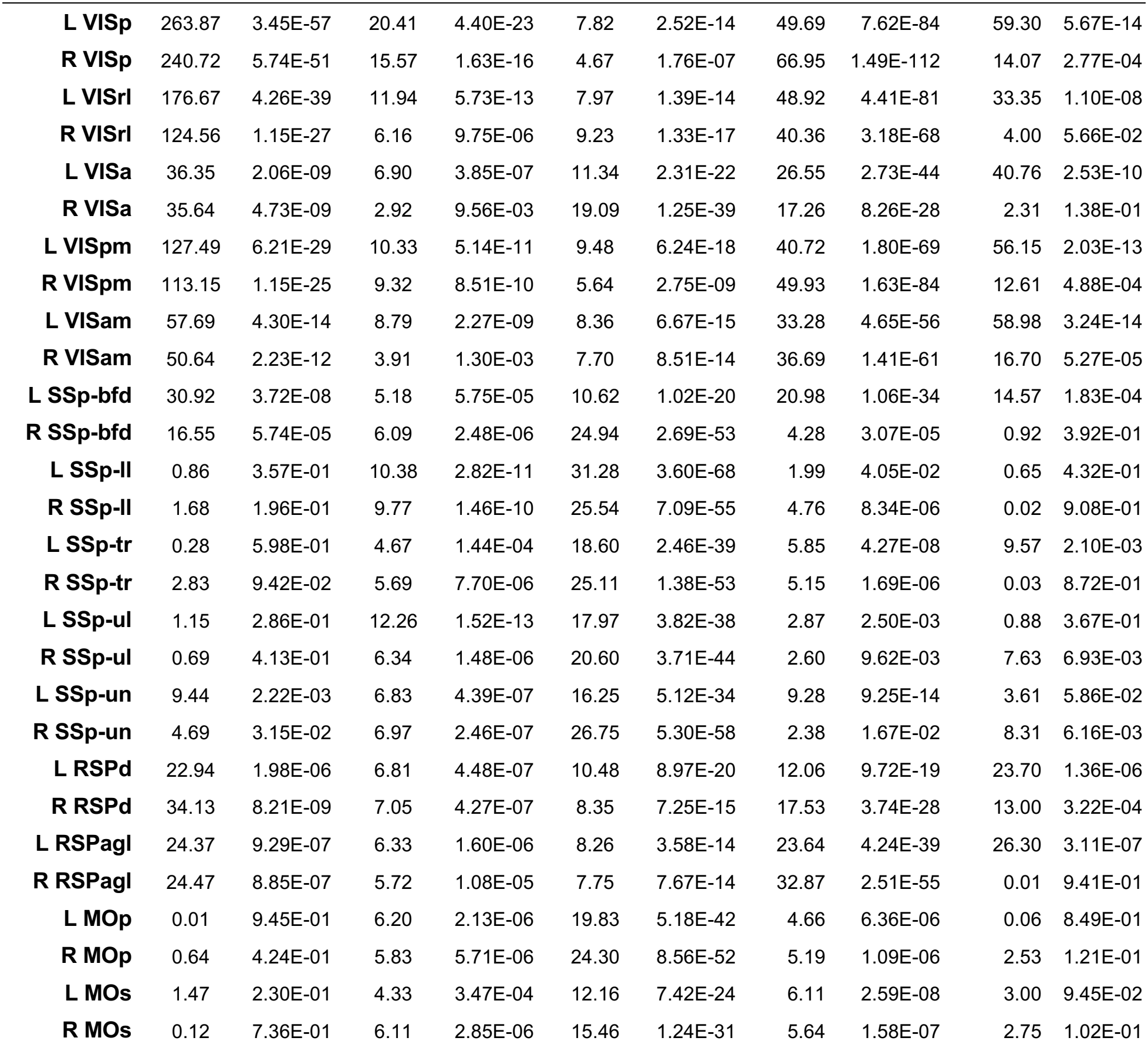
Statistics for the linear encoding model in Fig. 2, Extended Data Fig. 3. We used *F*-tests to compare the full model to nested models missing select predictors. *F* statistics and *P*-values are averages across five data splits.

| **Linear encoding model statistics** |  |  |  |  |  |  |  |  |
|---|---|---|---|---|---|---|---|---|
| **Pred.<br>removed** | **Coherence<br>(inst. + delayed)** |  | **Delayed<br>coherence** |  | **Test stimulus** |  | **Sample duration** |  |
|  | ***F*** | ***P*** **val.** | ***F*** | ***P*** **val.** | ***F*** | ***P*** **val.** | ***F*** | ***P*** **val.** |
| **Area** |  |  |  |  |  |  |  |  |
| **L VISp** | 22.82 | 5.03E-56 | 10.93 | 1.64E-21 | 2.42 | 8.99E-02 | 51.66 | 8.82E-60 |
| **R VISp** | 16.52 | 1.20E-37 | 8.16 | 5.65E-15 | 1.85 | 1.64E-01 | 44.89 | 4.90E-52 |
| **L VISrl** | 12.38 | 1.52E-28 | 6.47 | 2.30E-11 | 1.49 | 2.29E-01 | 38.15 | 7.47E-44 |
| **R VISrl** | 6.42 | 9.23E-13 | 4.13 | 2.01E-06 | 1.22 | 3.08E-01 | 29.62 | 1.18E-33 |
| **L VISa** | 6.72 | 1.41E-13 | 5.15 | 1.38E-08 | 3.80 | 2.41E-02 | 23.76 | 2.07E-27 |
| **R VISa** | 2.98 | 1.83E-04 | 2.31 | 6.51E-03 | 2.28 | 1.07E-01 | 16.58 | 1.37E-18 |
| **L VISpm** | 23.61 | 6.02E-58 | 13.51 | 2.26E-27 | 3.02 | 4.96E-02 | 43.22 | 1.57E-50 |
| **R VISpm** | 16.45 | 1.03E-37 | 9.45 | 8.86E-18 | 2.68 | 7.07E-02 | 40.71 | 3.31E-47 |
| **L VISam** | 13.01 | 5.07E-30 | 8.64 | 3.16E-16 | 2.91 | 5.59E-02 | 32.92 | 1.87E-38 |
| **R VISam** | 7.54 | 2.67E-15 | 5.25 | 1.21E-08 | 2.20 | 1.14E-01 | 28.17 | 1.49E-32 |
| **L SSp-bfd** | 1.73 | 4.84E-02 | 1.58 | 9.45E-02 | 2.34 | 1.06E-01 | 12.93 | 4.31E-14 |
| **R SSp-bfd** | 1.66 | 5.92E-02 | 1.88 | 3.36E-02 | 3.46 | 3.93E-02 | 8.29 | 9.03E-09 |
| **L SSp-ll** | 0.49 | 9.39E-01 | 0.53 | 8.95E-01 | 4.12 | 1.68E-02 | 5.84 | 5.63E-06 |
| **R SSp-ll** | 0.39 | 9.77E-01 | 0.43 | 9.53E-01 | 4.68 | 9.67E-03 | 7.03 | 2.38E-07 |
| **L SSp-tr** | 0.94 | 5.11E-01 | 0.97 | 4.71E-01 | 3.55 | 3.12E-02 | 8.36 | 6.06E-09 |
| **R SSp-tr** | 0.69 | 7.82E-01 | 0.71 | 7.40E-01 | 3.43 | 3.36E-02 | 6.58 | 7.71E-07 |
| **L SSp-ul** | 0.61 | 8.58E-01 | 0.53 | 8.92E-01 | 4.14 | 1.70E-02 | 6.49 | 1.00E-06 |
| **R SSp-ul** | 0.69 | 7.81E-01 | 0.71 | 7.38E-01 | 6.50 | 2.02E-03 | 7.40 | 1.12E-07 |
| **L SSp-un** | 0.75 | 7.23E-01 | 0.83 | 6.20E-01 | 3.69 | 2.87E-02 | 7.73 | 4.35E-08 |
| **R SSp-un** | 1.18 | 2.82E-01 | 1.25 | 2.44E-01 | 6.15 | 3.27E-03 | 6.02 | 3.88E-06 |
| **L RSPd** | 7.11 | 2.46E-14 | 4.74 | 1.23E-07 | 1.71 | 1.84E-01 | 25.34 | 1.69E-29 |
| **R RSPd** | 5.98 | 3.19E-11 | 3.86 | 1.15E-05 | 3.25 | 4.04E-02 | 27.75 | 1.63E-32 |
| **L RSPagl** | 9.82 | 2.19E-21 | 6.76 | 4.45E-12 | 2.99 | 5.27E-02 | 28.70 | 1.71E-33 |
| **R RSPagl** | 6.98 | 5.22E-14 | 4.73 | 1.40E-07 | 3.24 | 4.16E-02 | 26.23 | 2.40E-30 |
| **L MOp** | 1.22 | 2.53E-01 | 0.66 | 7.90E-01 | 2.11 | 1.22E-01 | 3.18 | 4.38E-03 |
| **R MOp** | 1.95 | 1.88E-02 | 1.02 | 4.32E-01 | 3.70 | 2.69E-02 | 2.95 | 7.57E-03 |
| **L MOs** | 1.09 | 3.60E-01 | 0.79 | 6.63E-01 | 1.61 | 2.01E-01 | 1.92 | 7.42E-02 |
| **R MOs** | 1.40 | 1.50E-01 | 0.96 | 4.83E-01 | 1.74 | 1.77E-01 | 2.31 | 3.15E-02 |
| **L VISp** | 12.73 | 4.63E-16 | 13.56 | 1.85E-06 | 11.66 | 9.06E-06 | 7.29 | 9.24E-06 |
| **R VISp** | 7.99 | 3.35E-09 | 9.50 | 1.09E-04 | 4.65 | 1.02E-02 | 9.12 | 2.83E-07 |
| **L VISrl** | 6.95 | 3.52E-08 | 2.63 | 7.98E-02 | 1.31 | 2.70E-01 | 5.36 | 2.80E-04 |
| **R VISrl** | 7.38 | 1.00E-08 | 0.16 | 8.56E-01 | 0.37 | 6.93E-01 | 4.43 | 1.62E-03 |
| **L VISa** | 10.21 | 2.25E-12 | 1.03 | 3.65E-01 | 0.27 | 7.66E-01 | 0.54 | 7.05E-01 |
| **R VISa** | 6.76 | 9.64E-08 | 0.07 | 9.33E-01 | 0.67 | 5.14E-01 | 0.54 | 7.05E-01 |
| **L VISpm** | 14.10 | 7.31E-18 | 8.10 | 4.02E-04 | 5.13 | 6.01E-03 | 1.13 | 3.44E-01 |
| **R VISpm** | 12.40 | 6.80E-15 | 5.72 | 3.84E-03 | 2.04 | 1.31E-01 | 3.35 | 1.02E-02 |
| **L VISam** | 11.88 | 2.06E-14 | 3.11 | 4.80E-02 | 0.33 | 7.24E-01 | 0.69 | 6.03E-01 |
| **R VISam** | 11.06 | 4.91E-13 | 1.28 | 2.86E-01 | 0.10 | 9.05E-01 | 0.97 | 4.30E-01 |
| **L SSp-bfd** | 4.01 | 2.33E-04 | 0.39 | 6.79E-01 | 0.95 | 3.92E-01 | 0.47 | 7.54E-01 |
| **R SSp-bfd** | 2.00 | 5.34E-02 | 4.71 | 1.06E-02 | 0.10 | 9.05E-01 | 0.69 | 5.97E-01 |
| **L SSp-ll** | 3.82 | 4.09E-04 | 0.83 | 4.39E-01 | 0.33 | 7.22E-01 | 0.34 | 8.51E-01 |
| **R SSp-ll** | 4.24 | 1.24E-04 | 0.28 | 7.56E-01 | 0.93 | 3.95E-01 | 0.79 | 5.37E-01 |
| **L SSp-tr** | 4.88 | 1.91E-05 | 0.95 | 3.93E-01 | 0.10 | 9.05E-01 | 0.25 | 9.10E-01 |
| **R SSp-tr** | 3.37 | 1.50E-03 | 0.56 | 5.76E-01 | 0.36 | 6.95E-01 | 0.97 | 4.26E-01 |
| **L SSp-ul** | 3.64 | 6.70E-04 | 3.19 | 4.26E-02 | 0.02 | 9.80E-01 | 0.64 | 6.36E-01 |
| **R SSp-ul** | 2.32 | 2.34E-02 | 3.99 | 1.99E-02 | 0.56 | 5.77E-01 | 1.00 | 4.08E-01 |
| **L SSp-un** | 3.48 | 1.05E-03 | 1.54 | 2.21E-01 | 0.78 | 4.67E-01 | 0.43 | 7.87E-01 |
| **R SSp-un** | 1.77 | 9.13E-02 | 7.45 | 6.84E-04 | 0.14 | 8.77E-01 | 0.52 | 7.20E-01 |
| **L RSPd** | 10.91 | 7.40E-13 | 1.96 | 1.46E-01 | 0.95 | 3.89E-01 | 0.89 | 4.70E-01 |
| **R RSPd** | 11.20 | 2.70E-13 | 0.96 | 3.94E-01 | 0.50 | 6.05E-01 | 0.52 | 7.20E-01 |
| **L RSPagl** | 10.44 | 1.94E-12 | 3.99 | 2.23E-02 | 0.43 | 6.55E-01 | 0.63 | 6.43E-01 |
| **R RSPagl** | 11.53 | 9.89E-14 | 5.79 | 4.25E-03 | 0.10 | 9.03E-01 | 0.35 | 8.45E-01 |
| **L MOp** | 3.81 | 4.13E-04 | 2.50 | 8.31E-02 | 0.07 | 9.30E-01 | 1.36 | 2.47E-01 |
| **R MOp** | 3.35 | 1.55E-03 | 5.14 | 6.37E-03 | 0.46 | 6.35E-01 | 1.94 | 1.04E-01 |
| **L MOs** | 3.06 | 3.48E-03 | 4.14 | 1.63E-02 | 0.36 | 6.95E-01 | 1.24 | 2.91E-01 |
| **R MOs** | 2.42 | 1.90E-02 | 6.22 | 2.20E-03 | 0.38 | 6.87E-01 | 1.61 | 1.69E-01 |
| **L VISp** | 263.87 | 3.45E-57 | 20.41 | 4.40E-23 | 7.82 | 2.52E-14 | 49.69 | 7.62E-84 | 59.30 | 5.67E-14 |
| **R VISp** | 240.72 | 5.74E-51 | 15.57 | 1.63E-16 | 4.67 | 1.76E-07 | 66.95 | 1.49E-112 | 14.07 | 2.77E-04 |
| **L VISrl** | 176.67 | 4.26E-39 | 11.94 | 5.73E-13 | 7.97 | 1.39E-14 | 48.92 | 4.41E-81 | 33.35 | 1.10E-08 |
| **R VISrl** | 124.56 | 1.15E-27 | 6.16 | 9.75E-06 | 9.23 | 1.33E-17 | 40.36 | 3.18E-68 | 4.00 | 5.66E-02 |
| **L VISa** | 36.35 | 2.06E-09 | 6.90 | 3.85E-07 | 11.34 | 2.31E-22 | 26.55 | 2.73E-44 | 40.76 | 2.53E-10 |
| **R VISa** | 35.64 | 4.73E-09 | 2.92 | 9.56E-03 | 19.09 | 1.25E-39 | 17.26 | 8.26E-28 | 2.31 | 1.38E-01 |
| **L VISpm** | 127.49 | 6.21E-29 | 10.33 | 5.14E-11 | 9.48 | 6.24E-18 | 40.72 | 1.80E-69 | 56.15 | 2.03E-13 |
| **R VISpm** | 113.15 | 1.15E-25 | 9.32 | 8.51E-10 | 5.64 | 2.75E-09 | 49.93 | 1.63E-84 | 12.61 | 4.88E-04 |
| **L VISam** | 57.69 | 4.30E-14 | 8.79 | 2.27E-09 | 8.36 | 6.67E-15 | 33.28 | 4.65E-56 | 58.98 | 3.24E-14 |
| **R VISam** | 50.64 | 2.23E-12 | 3.91 | 1.30E-03 | 7.70 | 8.51E-14 | 36.69 | 1.41E-61 | 16.70 | 5.27E-05 |
| **L SSp-bfd** | 30.92 | 3.72E-08 | 5.18 | 5.75E-05 | 10.62 | 1.02E-20 | 20.98 | 1.06E-34 | 14.57 | 1.83E-04 |
| **R SSp-bfd** | 16.55 | 5.74E-05 | 6.09 | 2.48E-06 | 24.94 | 2.69E-53 | 4.28 | 3.07E-05 | 0.92 | 3.92E-01 |
| **L SSp-ll** | 0.86 | 3.57E-01 | 10.38 | 2.82E-11 | 31.28 | 3.60E-68 | 1.99 | 4.05E-02 | 0.65 | 4.32E-01 |
| **R SSp-ll** | 1.68 | 1.96E-01 | 9.77 | 1.46E-10 | 25.54 | 7.09E-55 | 4.76 | 8.34E-06 | 0.02 | 9.08E-01 |
| **L SSp-tr** | 0.28 | 5.98E-01 | 4.67 | 1.44E-04 | 18.60 | 2.46E-39 | 5.85 | 4.27E-08 | 9.57 | 2.10E-03 |
| **R SSp-tr** | 2.83 | 9.42E-02 | 5.69 | 7.70E-06 | 25.11 | 1.38E-53 | 5.15 | 1.69E-06 | 0.03 | 8.72E-01 |
| **L SSp-ul** | 1.15 | 2.86E-01 | 12.26 | 1.52E-13 | 17.97 | 3.82E-38 | 2.87 | 2.50E-03 | 0.88 | 3.67E-01 |
| **R SSp-ul** | 0.69 | 4.13E-01 | 6.34 | 1.48E-06 | 20.60 | 3.71E-44 | 2.60 | 9.62E-03 | 7.63 | 6.93E-03 |
| **L SSp-un** | 9.44 | 2.22E-03 | 6.83 | 4.39E-07 | 16.25 | 5.12E-34 | 9.28 | 9.25E-14 | 3.61 | 5.86E-02 |
| **R SSp-un** | 4.69 | 3.15E-02 | 6.97 | 2.46E-07 | 26.75 | 5.30E-58 | 2.38 | 1.67E-02 | 8.31 | 6.16E-03 |
| **L RSPd** | 22.94 | 1.98E-06 | 6.81 | 4.48E-07 | 10.48 | 8.97E-20 | 12.06 | 9.72E-19 | 23.70 | 1.36E-06 |
| **R RSPd** | 34.13 | 8.21E-09 | 7.05 | 4.27E-07 | 8.35 | 7.25E-15 | 17.53 | 3.74E-28 | 13.00 | 3.22E-04 |
| **L RSPagl** | 24.37 | 9.29E-07 | 6.33 | 1.60E-06 | 8.26 | 3.58E-14 | 23.64 | 4.24E-39 | 26.30 | 3.11E-07 |
| **R RSPagl** | 24.47 | 8.85E-07 | 5.72 | 1.08E-05 | 7.75 | 7.67E-14 | 32.87 | 2.51E-55 | 0.01 | 9.41E-01 |
| **L MOp** | 0.01 | 9.45E-01 | 6.20 | 2.13E-06 | 19.83 | 5.18E-42 | 4.66 | 6.36E-06 | 0.06 | 8.49E-01 |
| **R MOp** | 0.64 | 4.24E-01 | 5.83 | 5.71E-06 | 24.30 | 8.56E-52 | 5.19 | 1.09E-06 | 2.53 | 1.21E-01 |
| **L MOs** | 1.47 | 2.30E-01 | 4.33 | 3.47E-04 | 12.16 | 7.42E-24 | 6.11 | 2.59E-08 | 3.00 | 9.45E-02 |
| **R MOs** | 0.12 | 7.36E-01 | 6.11 | 2.85E-06 | 15.46 | 1.24E-31 | 5.64 | 1.58E-07 | 2.75 | 1.02E-01 |

